# High-amplitude oscillatory events coordinate large-scale cortical interactions during decision-making and attention allocation

**DOI:** 10.1101/2025.11.21.689181

**Authors:** Marcus Siems, Yinan Cao, Tobias H. Donner, Konstantinos Tsetsos, Andreas K. Engel

**Author notes:** |aContact Information. Contributed equally (Konstantinos Tsetsos, Andreas K. Engel).

## Abstract

How do large-scale functional brain networks dynamically emerge to enable cognition? Correlated oscillations, a mechanism for inter-areal interactions, can be expressed as phase coherence or correlated amplitude fluctuations. While the functional role of phase coherence has been addressed in the past, it remains unknown how large-scale amplitude coupling dynamically supports the rapid network reconfigurations necessary for adaptive behavior. Here, we demonstrate that cognitive processing relies on the temporal alignment of brief, high-amplitude oscillatory events across distant cortical regions, rather than sustained coupling of oscillations. Capitalizing on the spatio-temporal resolution of magnetoencephalography (MEG) during a task probing attention and decision-making, we show that oscillatory event coincidences form functionally relevant, transient networks: First, coincidences of high-amplitude oscillatory events in the alpha and beta frequency range increase prior to correct decisions within a parieto-frontal network. Second, transient theta/alpha oscillatory event coincidences within a medial parietal, temporo-parietal junction and lateral prefrontal network track the dynamics of spontaneous, covert spatial attention reallocation. In contrast, transient aperiodic and sustained network components exhibit only weak functional modulations, revealing a spatial and spectral dissociation between transient and sustained amplitude coupling. Overall, our findings demonstrate that efficient cognition relies on large-scale interactions rapidly formed through co-occurring transient oscillatory events. We propose that the brain-wide orchestration of high-amplitude oscillatory events can provide a new framework for understanding how the brain dynamically integrates information to guide flexible behavior.

## Introduction

Cognition requires the exchange of information represented in distant areas of the brain. A central question in neuroscience is how local activity can be orchestrated between distant brain areas to integrate information necessary for cognition and goal-directed actions. A growing body of literature proposes neural oscillations, reflecting excitability fluctuations in local neuronal assemblies^1–6^, to provide a temporal mechanism supporting the information exchange. Correlations between local oscillations can provide the necessary flexibility to meet the cognitive and behavioral demands posed by dynamic environments^1,7–10^. Considerable work highlighted phase coherence between these oscillations as a mechanism for millisecond-precise inter-areal communication^7–9,11^. Yet, a second, distinct mode of intrinsic coupling exists: the co-fluctuation of local population amplitudes^7,8,12–14^, i.e., the temporally aligned strength of these excitability fluctuations. However, how and if amplitude coupling dynamically supports inter-areal communication during adaptive behavior remains elusive. Here, we propose that the coincidence of transient, high-amplitude oscillatory events across distant cortical regions can enable temporally precise large-scale interactions.

Currently, a considerable gap divides our understanding of these two large-scale intrinsic coupling modes^7,8^. Phase coupling is physiologically constrained by conduction delays and might enable or disable communication by aligning local excitability fluctuations and (oscillatory) spiking input^3,9–11^ supporting efficient perception^15–17^ and cognition^11,18–23^. In contrast, how and if the strength of these excitability fluctuations, as part of amplitude coupling, can impact neuronal information exchange remains unclear. While time-averaged amplitude coupling correlates with perception^16^, memory^24^, and decision-making^25^, and can dissociate from phase coupling during task performance^26,27^, rest^12^, and disease^28^, a major unresolved question remains: Can the coupling of local signal amplitudes support temporally precise, rapid network reconfigurations necessary for adaptive behavior?

Resolving these coupling modes dynamically over time^29^ can help to elucidate the role of precisely timed amplitude co-fluctuations in cognition. Emerging evidence shows that transient high-amplitude events locally interfuse sustained bandlimited neuronal activity^3,23,30–35^. Crucially, these high-amplitude events are heterogeneous, originating from either oscillatory or aperiodic transients^3,34,36^. High-amplitude oscillatory events^30,37,38^ reflect rhythmic excitability fluctuations that can occur without firing rate modulation and display narrow-band spectral power increases^4,23,39^. Moreover, aperiodic activity, another feature of healthy brain function, which is modulated during cognition^39–41^ and linked to time varying firing rate fluctuations^3,42^, can also display transient amplitude increases^36,37,39^. We reason that transient oscillatory events, aperiodic transients, and sustained low-amplitude activity support distinct information channels for neuronal interactions^3,42–44^. Hence, dissecting sustained amplitude coupling from transient oscillatory and aperiodic events should reveal distinct functional networks with dissociable relations to behavior. We hypothesized that the coincidence of high-amplitude oscillatory and aperiodic events^3,42^ enables the rapid cortical network reconfiguration required for flexible behavior. To test this hypothesis, we leveraged spatio-temporally precise magnetoencephalography (MEG) recordings to comprehensively unravel sustained and transient components of cortical activity while participants performed a challenging cognitive task^45^. Specifically, we identified functional networks defined through transient high-amplitude events coinciding in distant cortical areas and contrasted them against networks formed by sustained amplitude co-fluctuations. Assessing time-resolved signal rhythmicity^29,37^ enables us further to separate oscillatory and aperiodic high-amplitude event subtypes.

We show that coincident high-amplitude oscillatory events, but not sustained network activity, can predict decision-making performance and are modulated during attention allocation. Moreover, brain-wide networks defined by high-amplitude oscillatory event coincidences replicate functional networks previously identified via time-averaged amplitude co-fluctuations^12,13^. Overall, our findings demonstrate that cortical networks are dynamically shaped and regulated by transient high-amplitude events to support efficient cognition. We conclude that the temporally precise orchestration of coincident high-amplitude oscillatory events is a key signature of large-scale cortical interactions and can support efficient cognition.

## Results

### A framework for mapping cortical interactions via high-amplitude oscillatory events

Large-scale neuronal activity can be viewed as a vast space-time landscape of intensity fluctuations (Fig. 1). We hypothesized that transient high-amplitude oscillatory events play a distinct role in large-scale interactions indicating temporally precise orchestration between distant brain regions to form dynamic and flexible networks (Fig. 1a). Consequently, the temporal alignment of high-amplitude events across regions processing task-relevant information might directly impact behavioral outcomes, such as stimulus detection, when aligned to critical time points, e.g., the stimulus onset (Fig. 1b).

**Figure 1.**
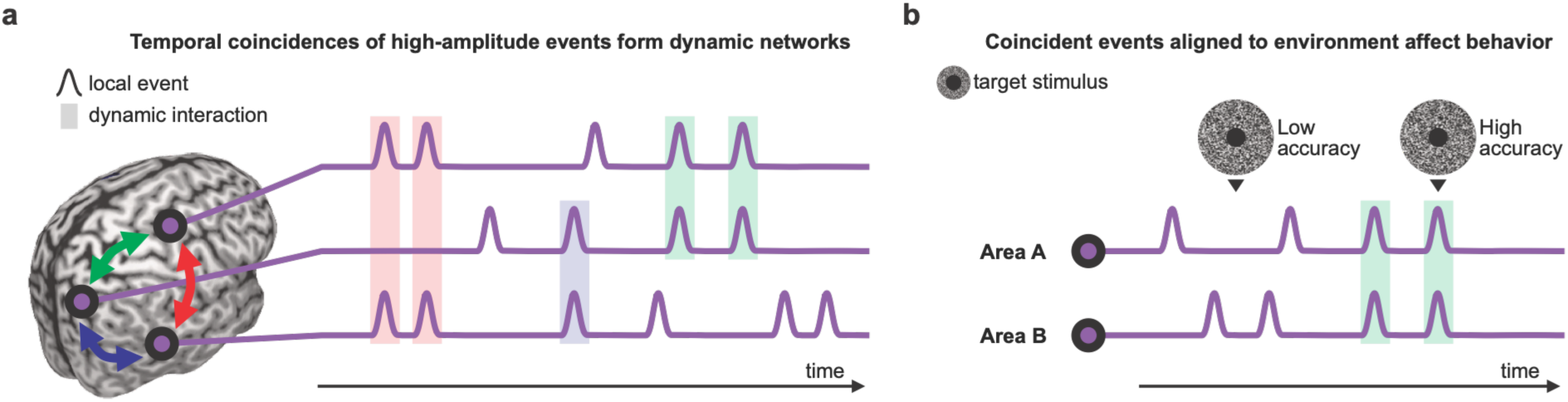
Schematic of high-amplitude oscillatory events orchestrating cortical activity for efficient cognition. **a** Temporal formation of dynamic networks (colored arrows) defined through coincident high-amplitude events (colored shading). **b** Dynamic networks forming by coincidence of events during critical periods of a task, for example the onset of a target stimulus, might impact performance in an attention task.

To test this framework, we recorded magnetoencephalography (MEG) from twenty healthy participants (9 female, age_mean_ = 28.05 years ± age_STD_ 4.6 years) performing a challenging three-alternative decision-making task^45^. Participants compared the contrast of three Gabor patches presented at peripheral locations equidistant from fixation and with variable task framing (median_accuracy_ = 77%, range_accuracy_ = 66–86%; see Methods). Harnessing (oscillatory) events, we established functional coupling based on the temporal coincidence of high-amplitude activity across distant cortical sources, termed oscillatory event coincidences (OECs). All amplitude coupling analyses were applied to pairwise orthogonalized signals to discount spurious coupling due to volume conduction^12,13^. Further, we separated oscillatory and aperiodic high-amplitude events using instantaneous frequency stability on source-reconstructed cortical activity over a broad range of frequencies (2–64 Hz; see Supplementary Text 1, Supplementary Fig. S1 and Methods).

We validated our novel approach in three complementary ways. First, we showed that local high-amplitude oscillatory event-rates align with previously reported^46^ cortical generators of oscillatory activity (see Supplementary Text 2 and Supplementary Fig. S2-S4). Second, time-averaged networks of amplitude coupling components displayed good reliability and, particularly, networks defined by transient OECs predicted previously reported amplitude coupling features (See Supplementary Text 3 and Supplementary Fig. S5-S10). Third, amplitude coupling network components were reliably and dynamically modulated during the task (Supplementary Text 4 and Supplementary Fig. S11-S14). In the following sections, we report how these distinct coupling modes relate to specific cognitive operations during task performance.

### Beta band oscillatory event coincidences predict correct decisions

We found that OEC networks were dynamically modulated during the task (Supplementary Text 4), and we tested whether transient amplitude co-fluctuation networks could predict decision behavior (Fig. 2). We first compared OEC dynamics between correct and error trials. Crucially, we matched the number of included (correct and error) trials by experimental conditions to control for task difficulty and sensory salience (see Methods). Averaging over all significant connections (n_connection_ = 104,196) and time points (4.45s time window, n_sample_ = 1,781) we identified significant coupling increases during correct trials spanning the high-alpha to beta frequency range (10–23 Hz), peaking in the low-beta range at 16 Hz (Fig. 2a; paired t-test(19), p_corrected_<0.05, cluster permutation corrected^47,48^ over sources, timepoints and frequencies).

**Figure 2.**
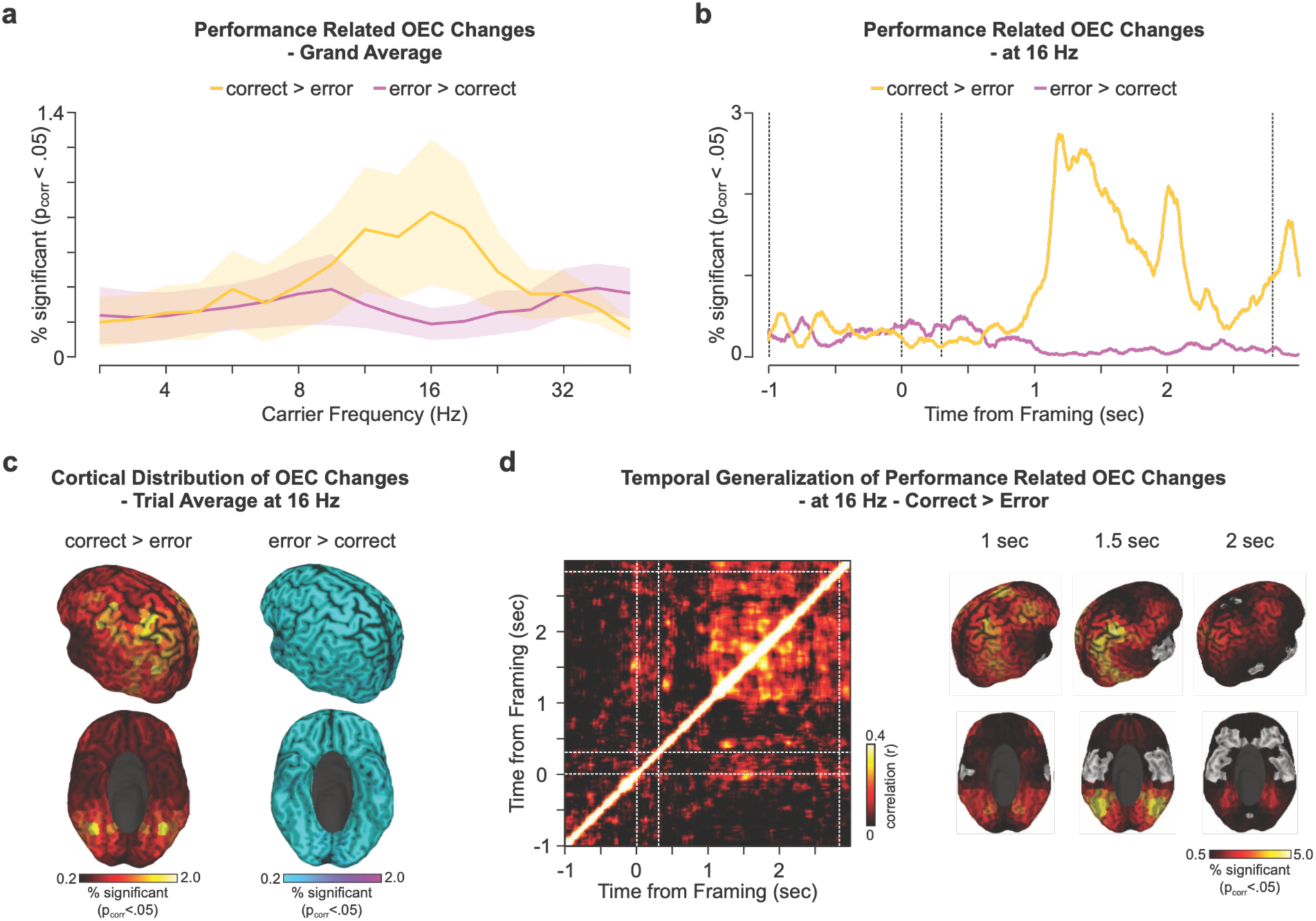
Oscillatory event coincidence rate at 16 Hz increases prior to correct decisions. **a** Spectrum of the total number of performance related (correct vs. error, two-sided paired t-test(19), cluster permutation corrected p<.05) connections per cortical source averaged over the full trial length (-1–2.95 s with respect to the framing cue). Orange and purple lines indicate whether OEC was increased during correct and error trials, respectively. Lines and shaded areas indicate the mean and standard deviation over cortical sources. **b** Time course of the total number of performance related connections at 16 Hz (two-sided paired t-test(19), cluster permutation corrected p<.05; color coded as above). Vertical dashed lines denote the stimulus onset, framing cue on- and offset and the stimulus offset (from left to right). **c** Cortical distribution of the number of performance related connections per source averaged over the full trial length at 16 Hz (two-sided paired t-test(19), cluster permutation corrected p<.05). **d** Temporal generalization of performance related coupling changes (correct > error) displayed through the cross-temporal correlation (left) of the cortical distribution of performance related coupling changes. Cortical distributions (right) show 0.5 s window averages for correct > error (one-sided paired t-test(19), cluster permutation corrected p<.025). Timing labels denote each window’s starting time with respect to the framing cue onset. The color-scale displays the number of significantly increased (correct > error) OEC connections per time window. The results were averaged over hemispheres.

We initially focused our analyses on this 16 Hz beta peak. We found that OEC rates increased primarily during the mid-decision phase (1–2 s post-framing) and immediately following stimulus offset. OEC rates increased particularly in the mid-decision phase around 1–2 seconds post-framing and right after stimulus offset (Fig. 2b). Correct choices were preceded by coupling increases within a medial and lateral parietal, posterior temporal, and lateral prefrontal network (Fig. 2c). This network appeared to be temporally stable over the decision phase (Fig. 2d and Supplementary Fig. S15). Conversely, a smaller subset of OEC connections showed relative coupling increases during error trials. These connections were restricted to the early visual and anterior temporal regions and during sensory and early decision phases (Fig. 2b,c). We further controlled for oculomotor activity and found no consistent overlap with decision-related functional networks (Supplementary Fig. S16). Together, these findings demonstrate that transient high-amplitude networks can be predictive of behavior: Beta-band OECs within an extended frontoparietal network were increased prior to successful decision-making.

### Transient and sustained amplitude coupling networks dissociate in decision-making

Having linked beta-band OECs to behavior, we mapped the behavioral relevance of OECs over the full frequency spectrum. Here, we tested whether partitioning amplitude coupling into transient (oscillatory and aperiodic) and sustained (event free) activity components can unravel distinct functional networks. Thus, we aimed at testing if any temporal, spectral or spatial features of decision-related network properties can be attributed uniquely to transient OECs.

Spectrally, performance modulated OEC increases (correct > error) predominantly spanned the beta range (Fig. 3a) and less pronounced decreases (error > correct) in the delta, alpha and gamma frequency ranges (Fig. 2a and Supplementary Fig. S17). Temporally, OEC increases occurred predominantly during the mid- and late decision phase (1–2.5 s from framing). On the other hand, OEC decreases peaked right after framing cue offset (0.3–1 s). Spatially, the grand average of OEC network changes (Fig. 3b) overlapped strongly with the network identified at 16 Hz (Fig. 2). Yet, within each frequency we identified dissociable networks reflecting performance (Supplementary Fig. S18): low-frequency OECs (4–8 Hz) increased in lateral parietal and lateral prefrontal areas; 11 Hz OECs recruited an extended parietal and posterior temporal network; and 22 Hz OECs increased in medial parietal and lateral prefrontal regions.

**Figure 3.**
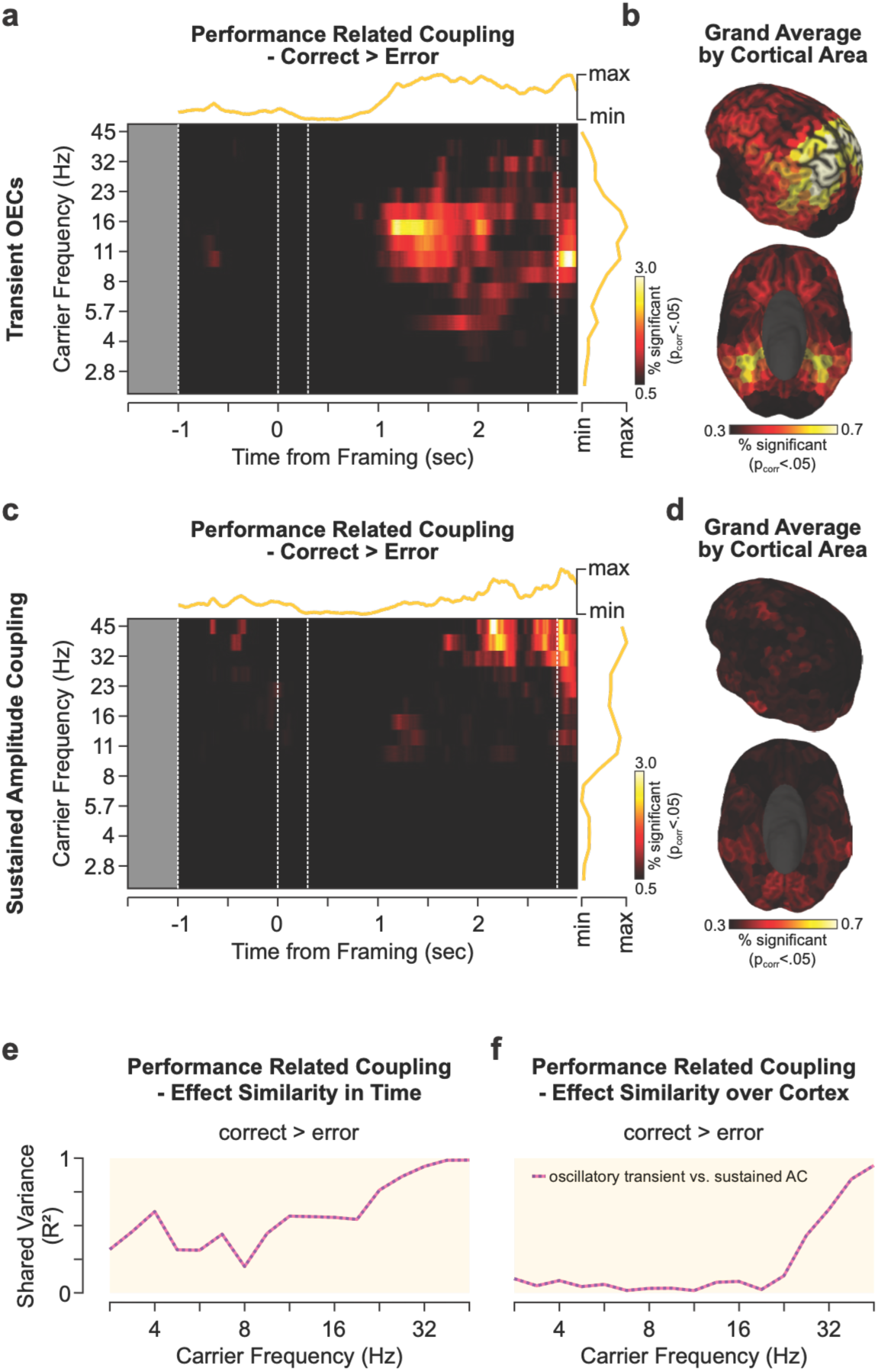
Spectral and spatial dissociation of performance related amplitude coupling networks. **a** Time and frequency resolved distribution of performance related (correct > error, one-sided t-test(19), cluster permutation corrected p<.05) transient OEC connections. Colored lines on top and to the right denote the average over frequencies and time, respectively. The pre-stimulus window (opaque, -1.5 to -1 s) was not tested for significance. **b** Average over the full time and frequency resolved distribution of performance related OEC modulations within each cortical source. **c-d** Same as **a-b** for sustained (event-free) amplitude coupling. **e-f** Pearson correlation of the behavioral coupling effects between transient oscillatory and sustained amplitude coupling (AC) comparing **e** the temporal (see Fig. 2a,c)

We next compared these findings to performance modulated amplitude coupling with (Supplementary Fig. S18-S20) and without high-amplitude events (Fig. 3c,d). Sustained (event-free) amplitude coupling revealed fewer, spectrally and spatially distinct task-related connections. Spectrally, sustained coupling decreased (error > correct) early in the trial at lower frequencies (4–8 Hz) but most prominently at higher frequencies (23–45 Hz; Figures S14 and S15). Spatially, performance modulated increases (correct > error) occurred in a fronto-parietal and early visual network and decreased in a medial parietal and ventral frontal network (Supplementary Fig. S18)

By comparison, transient aperiodic coincidences tracked correct choices within a parietal, temporal, and ventral-stream visual network that spatially overlapped with the transient oscillatory network (Figure 3b, Figure S18), with changes spectrally restricted to the high-beta to gamma range around stimulus offset.

Moreover, amplitude coupling (including events) performance effects mirrored a mixture of transient oscillatory and sustained amplitude network activity (Supplementary Fig. S18-S20). We, thus, systematically correlated the behavioral effects, both in time and over the cortex, between coupling components. The spatiotemporal correlation analyses revealed minimal network topography overlap between OECs and event-free coupling across all frequencies (Fig. 2e,f ; R^2^_cortex,2.8-45Hz_ = [<0.01, 0.04]) and only moderate temporal overlap of effects below the gamma range (R^2^_time,2.8-27Hz_ = [0.07, 0.48]). Furthermore, amplitude coupling effects (including events) could largely be explained through OECs for frequencies up to the beta ranges (R^2^_cortex,2.8-23Hz_ = [0.18, 0.76], R^2^_time,2.8-23Hz_ = [0.72, 0.97]). Yet, amplitude coupling (including events) networks aligned with sustained low-amplitude coupling at higher frequencies (Supplementary Fig. S18e,f; R^2^_cortex,27-45Hz_ = [0.43, 0.95], R^2^_time,27-45Hz_ = [0.87, 0.99]). Overall, these results demonstrate that decision-related networks dissociate temporally, spatially, and spectrally between transient and sustained coupling modes, with transient oscillatory coincidences accounting for the majority of interactions with choice behavior.

### Transient theta/alpha network activity tracks covert spatial attention allocation

Our analyses so far show dissociable links of transient and sustained components of dynamic amplitude coupling to decision-making behavior. We next investigated whether these coupling patterns map onto another facet of cognitive dynamics: the spontaneous allocation of covert spatial attention^49^. Previously, we could show that we can track the focus and strength of covert attention during the task^45^: Covert attention strength intrinsically fluctuates at 9–12 Hz, with attention reallocating between alternatives (“switch”) or refocusing on the same alternative (“stay”) at the trough and peak of this oscillation, respectively.

Here, we asked whether transient OECs and sustained amplitude coupling components are differentially modulated during attention allocation. We first quantified attention-triggered OECs at 10 Hz around the onset of attention reallocation refocus (Fig. 4a). Averaging over all connections revealed a significant increase of OECs during attention reallocation compared to refocusing (switch vs. stay, two-sided, paired t-test(19), p_FDR_<0.05; see also Supplementary Fig. S22 for associated resting-state networks). Extending this comparison across the full spectrum revealed coupling modulations spanning the theta, alpha, and beta bands (Fig. 4b). On the cortical level, between 3.5% (theta, 6 Hz) and 5% (alpha, 10 Hz) of all connections displayed increased transient OECs precisely during attention reallocation (per connection: paired t-test(19) of average OEC rate ±0.05 s from attention, switch vs. stay, p_corrected_<0.05, cluster permutation corrected over sources and frequencies). These switch-related OECs increases in the theta and alpha frequency range (Supplementary Fig. S23-S25) peaked in a network comprising ventral stream visual as well as medial parietal, temporo-parietal junction and lateral prefrontal areas (Fig. 4c). In contrast, stay-related coupling changes were sparse (Fig. 4c and Supplementary Fig. S25). We further controlled for oculomotor activity and found no consistent overlap with attention-modulated functional networks (Supplementary Fig. S18). Transient aperiodic networks showed no significant attentional modulation (Supplementary Fig. S26).

**Figure 4.**
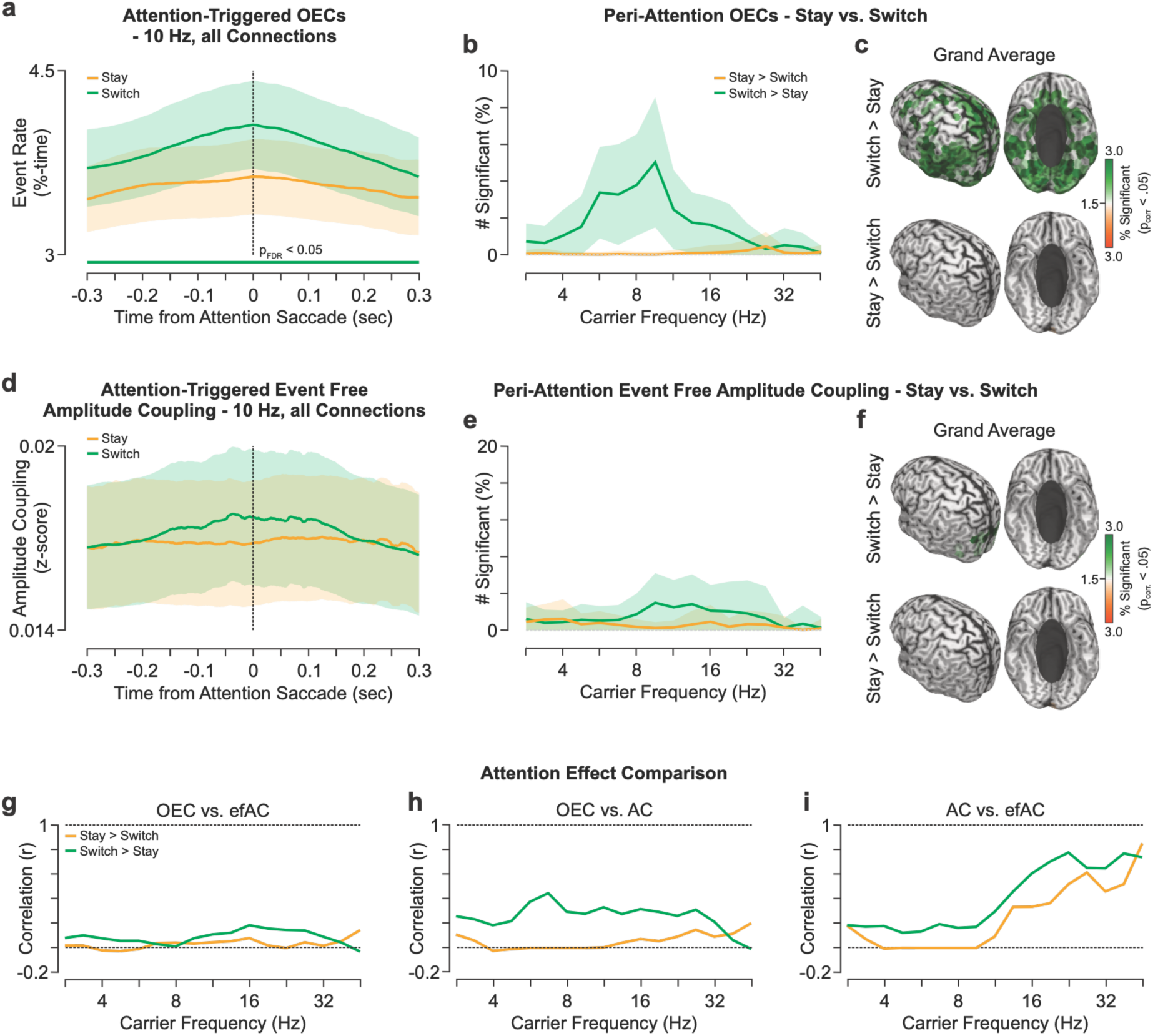
Amplitude coupling components dissociate during attention reallocation. **a** Attention-triggered OECs at 10 Hz separately for attention reallocation (switch, green) and refocus (stay, orange) averaged over all connections. Colored lines and shaded areas denote the mean and standard deviation over participants (n = 20). Thick lines denote the time of significant difference between switch and stay (two-sided paired t-test(19), pFDR<.05, FDR-corrected over time). **b** Number of OEC connections modulated by attention (paired two-sided t-test(19), cluster permutation corrected p<.05) at the time of attention allocation (averaged between -0.05 to 0.05 s) spectrally resolved between 2.8–45 Hz. Green and orange lines denote the mean proportion of connections per source increased during attention reallocation (switch) and refocusing (stay), respectively. Shaded areas denote the standard deviation over cortical sources. **c** Grand average of the proportion of attention modulated OEC connections from each cortical source averaged over all frequencies. **d-f** The same as **a-c** for sustained (event-free) amplitude coupling. **g-i** Correlation of attention modulation effects between coupling measures. The correlations were computed between frequency-specific cortical distributions of the number of switch- and stay-modulated connections (see Supplementary Fig. S24 and S25) for **g** OEC against event-free amplitude coupling (efAC), **h** OEC against full amplitude coupling including events (AC), and **i** between efAC and AC. Color-coded as above. and **f** spatial (Supplementary Fig. S18) distribution of effects. The values depict the shared variance (R^2^) within each frequency.

However, a very different relationship to attention emerged for sustained amplitude coupling (Fig. 4d-f). Amplitude coupling including high-amplitude events mirrored attention-modulated OECs spectrally and spatially (Fig. 4h and Supplementary Fig. S24). However, excluding high-amplitude events strongly reduced attention effects: Sustained coupling increased only in medial parietal regions during reallocation (Fig. 4f) and showed no significant attention modulation globally or within attention-related networks (Fig. 4d and Supplementary Fig. S22b). Across the full spectrum fewer connections modulated with attention (Fig. 4e; compare to Fig. 4b; see also Supplementary Fig. S23-S25). Again, this asymmetry between coupling components cannot be explained by differences in reliability or statistical power (Supplementary Fig. S6-S9).

The direct comparison of modulated attention networks confirmed this divergence (Fig. 4g-i, see Supplementary Fig. S23 & S24). Cortical network correlations during attention allocation revealed moderate overlap between OECs and amplitude coupling including events for frequencies up to the gamma range (Fig. 4h; switch; r_switch,2.8-32Hz_ = [0.17, 0.40]) and a strong correlations between standard and sustained (event-free) amplitude coupling at higher frequencies (Fig. 4i; r_switch,16-45Hz_ = [0.50, 0.81]; r_stay,16-45Hz_ = [0.19, 0.85]). Crucially, OEC and sustained amplitude coupling profiles remained weakly correlated across the entire spectrum with |r| < 0.25 (r_switch,2.8-45Hz_ = [-0.03, 0.24]; Fig. 4g). In sum, our findings demonstrate that transient theta and alpha OECs within a secondary visual to frontoparietal network track the rapid reallocation of covert attention and further underline a functional dissociation between transient and sustained cortical network dynamics.17

## Discussion

The exchange and integration of information over large distances in the brain is a prerequisite for flexible cognitive processing. Precise temporal relationships of neuronal activity reflected in phase coupling between distant areas have been associated with fostering and parsing this communication. However, little has been known about the functional role of correlated amplitude fluctuations of neural activity in long-distance communication and cognition. Utilizing non-invasive magnetoencephalography (MEG), we demonstrate that brief, high-amplitude oscillatory transients are a central hallmark of large-scale neuronal interactions during cognition. Specifically, the orchestration of these network-level transient oscillations shows to be predictive of successful decision-making behavior and dynamically tracks the rapid allocation of covert spatial attention. In contrast, sustained lower-amplitude co-fluctuations exhibit weaker task modulation and manifest in spatially and spectrally distinct networks. We, thus, conclude that our results provide empirical evidence that different components of large-scale amplitude co-fluctuations do not reflect a singular underlying network state, but might instead be supported by distinct neuronal circuit mechanisms.

At the physiological level, local activity strength and long-range amplitude co-fluctuations might arise from a combination of overlapping population-level mechanisms, including local neuronal synchronization, population firing rate modulations, and neuromodulatory excitability fluctuations^1,3,6,9,43,44^. Periodic excitability windows can temporally bias neuronal spiking without requiring macroscopic changes in population firing rates, leading to narrow-band oscillatory peaks in large-scale field potentials^1,6,7,14,39,40^. Less rhythmic population firing rate modulations can result in aperiodic spectral components in local field potentials^35,39–43,50^. Neuromodulatory inputs can broadly tune overall excitability and cellular firing readiness^6,43^. Critically, both local synchronization and aperiodic modulations can operate on fast timescales, generating transient, high-amplitude events^3,32–34,36^. We propose that tracking large-scale oscillatory and aperiodic events via signal rhythmivity^29,37^ offers a non-invasive readout of these micro-circuit operations: Macroscopic aperiodic events might reflect scale-free firing rate fluctuations at the cellular level^3,42^, whereas high-amplitude oscillatory events might capture the precise alignment of rhythmic excitability windows across distant regions^3,7,8^. However, future work utilizing invasive microscale recordings will be essential to directly validate how these distinct transient and sustained macroscopic signals map onto cellular firing dynamics and neuronal interactions.

Moreover, disentangling distinct components of amplitude coupling may enable the identification of temporally multiplexed channels for long-distance cortical information exchange. We found that sustained amplitude co-fluctuations as well as the transient coincidence of high-amplitude events, both oscillatory and aperiodic, form dissociable large-scale interactions, supporting the notion that they may represent functionally independent communication channels. Particularly, large-scale networks formed through transient oscillatory events are reliable, align with previously reported time-averaged amplitude coupling networks^12,13^, and offer independent insights into cortical interactions during cognition. As our results indicate that classical, time-averaged amplitude connectivity features were predominantly driven by transient high-amplitude events, prior findings across tasks and resting-state can be reevaluated with regard to the role of sparse, high-amplitude events. Our results help establishing these events as valuable non-invasive markers for intrinsic large-scale interactions, which can aid the study of healthy and atypical neuronal communication. In sum, rather than reflecting a continuous, uniform state, large-scale amplitude co-fluctuations might predominantly capture the precise orchestration between discrete, transient oscillatory network states.

Importantly, our multi-component dissection provides a framework to directly test previously reported, competing theories of large-scale neuronal communication. When local high-amplitude oscillatory events coincide across distant brain regions they also represent windows of heightened coherent activity^3,9,10,30,31^. Under the ‘Communication through Coherence’ (CTC) model of large-scale interactions these transient periods can align sender and receiver at excitable phases to enable the direct information exchange^20,50^. Computational models have highlighted strong oscillatory activity as a prerequisite for this phase-alignment^10^. This suggests that the inter-areal phase relationship within each discrete event coincidence might determine whether downstream information is effectively routed or suppressed^6,9,19,49,50^.

Our findings stipulate distinct empirical predictions for alternative models like ‘Coherence through Communication’ (CTC*) and ‘Communication through Resonance’ (CTR)^3^. According to CTC*, coherence is a statistical byproduct of population interactions that occur through correlated output of the sending area with its own input to the receiving area. Here, the inter-areal phase difference is structurally fixed via axonal conduction delays and synaptic time constants. Consequently, the phase relationship during high-amplitude oscillatory events should remain stable regardless of the cognitive state. In CTR, intrinsic structural properties affect the receiving population such that it selectively resonates with specific input frequencies. Consequently, the instantaneous frequency of the sending unit should exhibit high variability, i.e., low rhythmicity, prior to establishing a transient interaction, marking the transition towards the resonance frequency^3^. Overall, fully characterizing how distinct amplitude coupling components interact with phase coupling remains pivotal to understanding the mechanisms of flexible information _routing3,9,23,35._

Moreover, network activity is dynamically modulated during decision-making, our results provide empirical evidence that transient oscillatory events can act as distinct vehicles for the rapid, flexible network reconfigurations required for adaptive and goal-oriented behavior. During correct choices, transient oscillatory coupling selectively increased between medial and posterior parietal areas and the lateral prefrontal cortex, recruiting a network implicated in output preparation and sensory evidence comparison^51^. In contrast, sustained amplitude coupling was evident in the lateral parietal and medial prefrontal cortex, i.e., regions typically associated with integrative functions like error monitoring and evidence accumulation^52^.

Beyond decision processes, the precise alignment of theta and alpha OECs with spontaneous attention allocation shows that transient oscillatory coupling can reveal the mechanisms underlying the dynamics of covert behavior. During attention reallocation, transient theta/alpha networks comprising early visual, posterior parietal, and lateral frontal areas were modulated^45,53^. In contrast, maintaining the focus of attention (refocusing) was more strongly associated with sustained amplitude coupling in prefrontal areas. Together, these findings suggest that cognitive processes involving either change detection or steady-state information integration rely on distinct communication channels utilizing transient or sustained amplitude dynamics, respectively. In sum, our findings support the notion of a distinct relationship of cognitive processes involving either change detection or information integration with neuronal communication utilizing transient or sustained activity, respectively.

Several avenues remain for future research to address current limitations. First, the explicit mapping between cognition, macro-scale signal dynamics and meso- to micro-scale circuit-level mechanisms requires the direct investigation via combined intracranial and non-invasive recordings. Second, pharmacological interventions targeting arousal and the excitation/inhibition ratio can be leveraged to non-invasively probe these circuit-level mechanisms^37,39–41,43^ and could be used to elucidate malfunctioning coupling in the neurodiverse brain. Third, non-invasive electrical stimulation, i.e. multi-site amplitude-modulated transcranial alternating-current stimulation (AM-tACS) ^54^, can help to establish causal links between intrinsic coupling modes and cognition^8^. The amplitude modulation can induce transient peaks in oscillatory network activity to test the behavioral relevance of coincident oscillatory events.

In sum, our study demonstrates that large-scale amplitude coupling is a dynamically rich, functionally relevant intrinsic coupling mode that is dominated by the coincidence of transient, high-amplitude oscillatory events. Averaging functional coupling over extended time periods might obscure this highly variable interaction mode. Shifting the conceptual focus from continuous, sustained synchronization to a spatiotemporal landscape of transient events provides a new framework for understanding how the brain dynamically integrates information to guide behavior. By isolating oscillatory event coincidences, we reliably reconstructed large-scale amplitude coupling networks in the delta, theta, alpha and beta frequency ranges with only a fraction of the data. In contrast, sustained amplitude coupling, as well as aperiodic transients, captured reliable yet separate network features. Thus, sustained and transient coupling components might cater to qualitatively distinct functions and information processing time scales. Our results demonstrate that transient oscillatory networks are a key signature of inter-areal interactions and we argue that efficient cognition relies on information channels formed through co-occurring transient events.

## Methods

### Participants and experimental procedure

#### Participants

The task and participant group has been described in detail in^45^. In short, we tested twenty right-handed participants (9 female, age_mean_ = 28.05 years ± age_STD_ 4.6 years) in a decision-making task. The experiment was approved by the ethics committee of the Hamburg Medical Association and conducted in accordance with the Declaration of Helsinki. All participants gave their written informed consent and received monetary compensation for their participation.

#### Decision-making task

The main task was a three-alternative forced-choice protracted decision-making task with variable task framing. During central fixation three peripheral Gabor patches (2.2° eccentricity, 1.6° stimulus diameter) of varying contrast levels and orientations appeared simultaneously on screen (at the top and 120° left and right of it). The three stimuli were drawn from a set of five linearly spaced contrast levels (10 possible combinations, i.e., stimulus conditions). We used for each participant a staircase procedure to find the threshold physical distance between neighboring contrast levels and to control the task difficulty. The Gabor patch orientations were drawn from a set of three equally distant orientations with either all three being identical or different and pooled our analyses over orientation conditions. We did not utilize the orientation feature in the presented analyses.

Each trial started with the presentation of the central fixation cross. Participants were instructed to keep fixation and blink as little as possible during the trial. After 1 second the three Gabor patches appeared on screen and remained unaltered for 3.8 seconds. The task framing, i.e., if participants had to choose the highest (“Hig”) or lowest (“Low”) contrast, was centrally cued between 1–1.3 seconds into the stimulus presentation and randomized between trials. The high contrast stimulus represents the highest value in high framing trials, but the lowest value in low framing trials. The task framing divides the stimulus presentation period into the sensory- (pre- framing, -1–0 seconds from framing cue) and decision-phase (post-framing, 0.3–2.8 seconds) with identical visual stimulation but distinct cognitive demands. After the 3.8 second stimulus presentation, the Gabor patches turned off and the response cue (fixation cross rotated 45°) started the maximum 2 seconds response window. Participants indicated their decision with a left thumb, right thumb or foot button press for the left, right or top alternative, respectively. We randomized the foot (left or right) between participants. We individually adjusted the height and location of the pedal for each participant using Styrofoam such that the position was comfortable and a response could be triggered with a small flexion of the porcellus fori. We gave binary feedback after each trial (fixation cross: green for correct, red incorrect). We fully randomized task framing, contrast levels, Gabor orientation and stimulus locations, yielding 1080 unique trials. Every participant concluded two full randomization runs. We blocked the trials into random 120- trial sets (n_block_ = 18). Participants chose the correct contrast on average in 77% of all trials (range = [66% 86%]). For the analyses comparing decision-making performance (correct vs. error) we matched the trials included for both performance levels by stimulus conditions and framing cue to account for difficulty and salience effects.

#### Stimulus localizer task

Prior to each block of the decision-making task the participants concluded a localizer block: 90 trials of consecutively presented single Gabor stimuli in one of the three locations at full contrast for 0.35 seconds with 0.5 seconds inter-trial intervals. The stimuli matched the decision-making task in size and eccentricity. Participants were instructed to ignore the peripheral stimuli, fixate at the center and detect a flickering of the central fixation cross that could occur randomly at each stimulus. In total, a flicker occurred 9 times per block, and participants indicated the detection with a button press. In each block we fully randomized the orientation and the position over the three locations. Each stimulus combination was repeated ten times paired once with a central fixation flickering. We used the localizer task to decode the visual representations of all three stimuli during the decision-making task.

### Functional data acquisition and preprocessing

#### Data acquisition

During the experiment the participants were sitting upright in a whole-head magnetoencephalography scanner (MEG; 275 axial gradiometer sensors, CTF Systems Inc.) in a magnetically shielded room. The data was acquired at a sampling rate of 1200 Hz. Each participant’s head position was tracked online using three head localization coils (nasion, left/right preauricular points). Additionally, we recorded a 2-channel bipolar electro-oculogram (horizontal and vertical EOG) and a single channel bipolar electro-cardiogram (ECG) to control for cardiovascular and ocular signal artifacts. Eye-movements and pupil diameter were tracked using an MEG-compatible Eyelink 1000 system (SR Research). We further collected T1-weighted structural magnetic resonance images (MRI; sagittal MP-RAGE) to reconstruct individualized high-resolution head models.

#### Preprocessing

The detailed MEG preprocessing pipeline can be found in^45^. We band- pass filtered the continuous MEG data segments between 0.2 and 200 Hz (4^th^ order Butterworth filter) and down-sampled it to 400 Hz. We filtered the line noise with a notch filter at 50 Hz and its first 6 harmonics (1 Hz stop-band filter width). Next, we implemented a two-stage procedure for artifact rejection^55^ applying filtering the signal into two distinct frequency ranges (low- from 0.2 to 30 Hz; high-range from 30 to 200 Hz), separately computed independent component analyses on each range and rejected artifact components based on their topology, power-spectra and time-courses^56^.

### Spectral analysis and source reconstruction

#### Spectral analysis

We generated time-frequency resolved MEG signals using Morlet’s wavelets^55^ (wavelet bandwidth at 0.5 octaves; f/σ_f_ = 5.83; kernel width covered ± 3σ_t_; σ_f_ and σ_t_ corresponds to standard deviation in frequency and time domain). We derived complex spectral estimates for frequencies between 2 to 90.5 Hz in quarter octave steps (2^1^ to 2^6.5^ Hz).

#### Source projection

We generated single-shell boundary-element method models (BEM) based on individual structural MRIs. The physical forward model (leadfields) for 457 equally spaced cortical sources (∼1.2 cm distance, at 0.7 cm depth below pial surface) was computed using FieldTrip^57^. We applied dynamic imaging of coherent sources (DICS) linear beamforming to estimate the source-level neuronal activity. For seed-based analyses we used cortical source positions in the left auditory cortex (lAC, [-54, -22, 10]), left somatosensory cortex (lSSC, [-42, - 26, 54]), medial prefrontal cortex (MPFC, [-3, 39, -2])^12^, posterior medial temporal lobe (MT, [±39, -83, 15]), medial parietal cortex ([0, -64, 67]), lateral parietal cortex ([±41, -68, 44]), and lateral prefrontal cortex (PFC, [55, 13, 35]), all part of the source-model and in MNI-coordinates.

### Defining transient high-amplitude events

We defined transient high-amplitude events as episodes of bandlimited high-amplitude activity. Here, we z-scored the squared amplitude, i.e., signal power, of the complex time-frequency resolved data over the full experiment within each cortical source. Prior to z-scoring, we subtracted the trial-averaged activity. Within each source and frequency, we recorded high-amplitude events as time points with a z-score larger 3.

#### Oscillatory vs. aperiodic events

To identify if a sample within a high-amplitude event is oscillatory or not, we leveraged the phase-progression stability of band-limited oscillatory activity^37^. A pure oscillation exhibits a progression through the phase cycle with uniform speed, i.e., a stable instantaneous frequency (see Supplementary Fig. S1). A filtered aperiodic signal on the other hand will display a more variable instantaneous frequency distribution through intrinsic frequency modulation. First, we computed for each time point ***t*** and source ***i*** the cross-spectrum *c* between the signal ***x_t_*** with its past ***x_t-1_***.

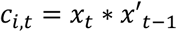

Where ***x’*** denotes the complex conjugate of the signal. We then derived the first-order temporal derivative of the instantaneous phase ***Δφ*** as the inverse tangent of the imaginary, divided by the real part of the cross-spectrum^29^.

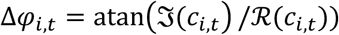

The absolute value ***||*** of the instantaneous phase derivative describes the phase progression stability over time. The instantaneous frequency ***f_i,t_*** can then be derived.

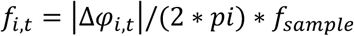

Where ***f_sample_*** denotes the sampling frequency of the signal. Finally, we defined oscillatory time points as neighboring samples displaying the lowest first-order temporal derivative of the instantaneous frequency (IFD), i.e., moments of the most stable phase progression, by

thresholding.

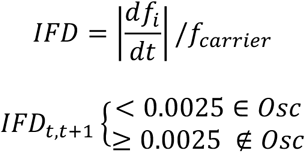

In other words, if the instantaneous frequency difference between neighboring samples, ***t*** and ***t+1***, is smaller than 0.25% of the carrier frequency ***f_carrier_*** we considered time point ***t*** to be most likely oscillatory. For a pure sinusoid the IFD will be equal to zero (see Supplementary Fig. S1). Overall, the approach needs only three samples of the signal to generate a proxy of oscillatory activity, i.e., at a sampling rate of 400 Hz only 0.0075 seconds. The threshold parameter was derived as the approximate median 25^th^ percentile of the IFD over the cortex at 11 Hz (2^3.5^ Hz). Thus, at 11 Hz approximately 25% of time points for each source are set to be most likely oscillatory. We selected the alpha range as a reference frequency due to the large body of literature identifying oscillatory alpha activity^37,39,44^. We then applied the same threshold to all analyzed carrier frequencies. The threshold scales well with carrier frequency because the constructed Morlet wavelets filter widths are adaptive.

Lastly, we defined oscillatory and aperiodic high-amplitude events through the co-occurrence or not co-occurrence with a stable instantaneous frequency, respectively.

### Analyzing event dynamics

#### Event rate and duration

We computed the local oscillatory event rate as the percentage of time points that an oscillatory event occurred at a given source and frequency. Under random conditions, i.e., no oscillatory activity, the minimum expected event rate can be derived from multiplying the probabilities of the signal crossing the amplitude and the IFD threshold (for frequencies: median_2-64Hz_ = 0.26%, range_2-64Hz_ = 0.03–0.34%). The relative event rate is the percentage of oscillatory events among all recorded high-amplitude events. Under random conditions the average relative oscillatory event rate would converge towards the probability of crossing the IFD threshold parameter (for frequencies: median_2-64Hz_ = 29.3%, range_2-64Hz_ = 2.7– 35.2%). We applied fixed thresholds to explicitly address potential differences in the frequency-specific amplitude (z_threshold_ > 3) and instantaneous frequency (Δf_threshold_ < 0.25%) distributions. The duration describes the median length of each high-amplitude event. Due to adaptive filter width of the applied Morlet wavelets the amplitude modulation can only depict a smaller range for lower frequencies and durations, measured in units of time, will be longer. We thus also computed the duration in units of cycle length by multiplying with the carrier frequency.

### Functional coupling analyses

#### Amplitude coupling

We applied two different approaches to quantify the statistical relationships between the signal strength of distant cortical sources: amplitude coupling and event coincidences. For the amplitude coupling we pairwise multiplied (dot product) z-scored power envelope (squared amplitude) signals between a seed and a target region. These seed-target cofluctuations were either averaged over time, yielding the correlation coefficient (time-averaged coupling analysis) or used to describe the temporal dynamics of amplitude coupling^29^. To discount effects of volume conduction, we orthogonalized the target-onto the seed signal prior to squaring and z-scoring the power envelopes^13^. The z-scoring was conducted over the full dataset within each cortical source and frequency. For the full correlation matrices (source x source) we applied the orthogonalization in both direction with signals x and y as the seed and target and vice versa.

#### Coupling by event coincidence

We quantified coincidences of high-amplitude oscillatory events (OECs) between distant cortical sources as the “logical and” operation for the occurrence of high-amplitude events and stable instantaneous frequency in both the seed and target signals at the same time. Aperiodic events on the other hand showed high amplitude without IF stability in both sources. Hereby, the definition of high-amplitude events was based on the same orthogonalized z-scored power envelopes as the amplitude coupling. Coincidence under random conditions was estimated by multiplying the respective local event rates from seed and orthogonalized target, i.e., co-occurring events without temporal relationships between seed and target. The oscillatory (and aperiodic) coincidence time courses thus are binarized signals with zeros and ones indicating no coincidence and coincidence, respectively. Again, averaging these signals over time yields the coincidence rate for a specific connection (time-averaged coupling analysis), while the binary time courses were used to describe dynamic coupling with respect to external and internal events. Additionally, we computed event free amplitude coupling by excluding all local event time points from the seed-target power cofluctuations, i.e., we set high-amplitude event time points to nan (not a number).

#### Statistical testing

We tested for statistical coupling differences between conditions using parametric paired t-tests (df = 19). We controlled for multiple comparisons using either false-discovery-rate correction^58^ (FDR-correction) or cluster-based permutation statistics^48^ using the fieldtrip toolbox^47^. For the cluster permutation analyses we formed cluster cortical neighborhoods (hexagonal mesh of cortical sources^59^), timepoints and frequencies for performance modulated coupling during decision-making (correct vs. error; n_permutations_ = 200) and without the time dimension for attention modulated coupling (switch vs. stay; n_permutations_ = 500). We quantified the cluster mass (p < 0.05) through the summed t-scores (‘maxsum’ criterion) within each cluster.

### Data simulation

To highlight the distinct features of oscillatory and aperiodic events we simulated 5 second traces of random data *x* with a pink noise spectrum (power ∼ 1/f; n_simulation_ = 100,000). We split each data trace into five 1-second windows (I-V). We multiplied the data in window II by 3 (*3x*) to increase broadband (aperiodic) power. We further multiplied window IV by 0.8 and added a constant 10 Hz sine wave *y* with an amplitude of 0.8 (*0.8x + 0.8y*), to increase oscillatory activity and bandlimited power. The modulation parameters were chosen to i) approximately match the broadband power between the random data segments (I, III, V) and the sine modulated window IV as well as ii) approximately matching bandlimited power between 8–12 Hz for both modulated windows (II and IV). We estimated the broadband power within each window as the integral of the windowed Fourier transform. We applied the same Morlet wavelet transformation as outlined for the empirical data with a center frequency at 10 Hz and estimated the bandlimited power from the broadband Fourier transform solution between 8–12 Hz. We applied the event detection algorithm on the bandlimited full data trace while z-scoring the data according to the mean and standard deviation from windows I, III, and V for each simulation repetition. The IF stability threshold was set to the 10% percentile over windows I, III, and V.

### Encoding the focus of covert attention

We used a stimulus localizer task to reconstruct the competition of stimulus representations during the decision-making task and thereby track the focus and strength of attention. Here, we implemented an inverted encoding model^45^. The encoding model assumes that the sum of abstract neuronal populations, i.e., information channels responsive to a given stimulus angle, is represented in the multivariate sensor-level MEG activity. Activity and tuning for an information channel can thus be estimated as the weighted sum of this MEG activity plus noise. The activity derived from inverting the model for MEG activity in the decision-making task then denotes momentary changes of the visual representations of the simultaneously presented stimuli.

#### Training the encoding model

We trained the encoding model on the retinotopic stimulus localizer task. During the localizer task only one stimulus out of the full set has been visible on screen. Hence, during the localizer both neuronal activity and the to be represented visual stimulus were known variables. We used bell-shaped tuning-curves (half-wave rectified sinusoidal) as design matrix to identify a linear weight-matrix relating neuronal activity and the presented stimuli^45^. We further used multivariate noise normalization on the weight-matrix to address correlated sensor activity, i.e., volume conduction, and improve reliability.

#### Applying inverted model on task data

With the optimized weight-matrix we can estimate the latent visual representation of the stimuli during the multi-alternative decision-making task. We inverted the model, i.e., the weight-matrix, and multiplied the MEG activity during the decision-making task. The solution of this operation yielded the representational activity over visual stimuli for each time point during the task. The strength of the representations is a proxy of covert visual attention towards each target on screen.

#### Reconstruction of attention focus

We used the vector sum of the three representational activities as a handle on the attention focus^45^. The vector orientations were set to 0°, 120° and 240° for the top, left and right stimulus, respectively and the representational activity corresponded to the length at each orientation. The reconstructed attentional vector can illustrate two features of visuo-spatial attention: First, the overall attention/alertness^60^ paid to the stimuli and second, the main target of information sampling. Thus, we considered the length of the vector as a proxy for overall attention and the direction, i.e., the angular similarity to the three alternatives, as the most likely target of covert spatial attention at each time point. Importantly, this quantification of covert attention cannot be solely explained by fixational eye-movements or (micro-)saccades^45^.

*Model training parameter.* We optimized the decoding performance within the localizer task by varying the training time point with respect to stimulus onset. Applying leave-one-block-out cross-validation (n_block_ = 18 per participant) we selected the time-window from 0.14–0.17 seconds after stimulus onset as training epoch^45^. For the model training we excluded stimulus localizer trials displaying (micro-)saccadic activity within the first 0.2 seconds after stimulus onset. On average 3% of all localizer trials per participant (mean = 3.03%, median = 1.82% range = 0.50% - 10.68%) were excluded.

#### Defining moments of attention reallocation and refocus

We leveraged the temporal dynamics of the attention vector to define moments when attention shifted from one alternative to another. The momentary focus of spatial attention was assessed via the angular similarity between the attention vector and the vector orientations corresponding to the three stimuli. At every moment the vector can point close to one decision alternative or in between two options. We defined a threshold to identify time points where attention is focused at one alternative as the 90^th^ percentile of the angular similarity distribution^45^. However, the vector is oriented in circular space and will even under random conditions, i.e., without visual stimulation or participant’s alertness, point towards one of the three stimuli. We thus applied a second threshold on the vector length (within trial 90^th^ percentile). An attentional saccade is only recorded when both thresholds, i.e., orientation and length of the vector, are crossed at the same time. We only recorded the first sample when both thresholds were crossed and excluded the following 0.025 seconds to not count individual events twice. We previously assessed the effects of different thresholds and concluded that the selected parameter range is sensible and does not qualitatively affect the conclusions on spatial attention (dynamics)^45^. Using this approach we identified 17,696 attention saccades per participant (median; range 12,377 to 28,909). We further tracked first-order sequences: Does an attention saccade focus on the same stimulus as the previous one (“stay”, refocus) or shift between alternatives (“switch”, reallocate). These sequences appear to be related to distinct attention functions and neuronal correlates^45^.

#### Attention-related high-amplitude event dynamics

To assess the neuronal activity related to attention reallocation and refocus we computed the average event coincidence rate and amplitude coupling dynamics between 0.3 seconds before and after each attention reallocation/refocus. Importantly, we matched the included switch and stay by trial timing because the trial dynamics between both types vary^45^. Thus, we effectively included around 5,500 saccades per type and participant (median = 5,627, range = 3,824–10,771). We statistically tested neuronal activity between attention types, i.e., switch versus stay, applying paired t-tests of the attention-averaged activity for each sample within the attention-triggered time window (n_sample_ = 241). We applied false-discovery rate correction (FDR^58^) within each window. For analyses summarizing the activity directly at the events (for example Fig. 4) we first averaged the data between -0.05–0.05 seconds (relative to the event) before we conducted paired t-tests on the time-averaged activity. For visualization we smoothed the peri-event activity by a 0.025s boxcar kernel.

## Data and code availability

The processed data generated in this study will be made publicly available upon publication. The code for reproducing the main results and figures will be made publicly available upon publication.

## Acknowledgements

This work was supported by grants from the European Research Council, project INFOSAMPLE, ERC-2018-StG-802905 (awarded to K.T.), and project cICMs, ERC-2022-AdG-101097402 (awarded to A.K.E.). Views and opinions expressed in this paper are those of the authors only and do not necessarily reflect those of the European Union or the European Research Council. Neither the European Union nor the granting authority can be held responsible for them. We thank Maryam Tohidi-Moghaddam for her help with the task design and data collection.

## Author contributions

M.S., K.T. and Y.C. contributed to the experimental protocol. M.S. and Y.C. collected the data.

M.S. conceptualized the functional coupling analysis. M.S. & Y.C. did the formal data analysis and M.S. visualized the results. M.S., T.H.D. & A.K.E. wrote the initial draft, all authors contributed to editing the manuscript. K.T. and A.K.E. acquired the funding for the study.

