## Supplementary Information for "High-amplitude oscillatory events coordinate large-scale cortical interactions during decision-making and attention allocation"

[*] Corresponding author

[+] Contributed equally

**Contact Information**

### Supplementary Information

#### Supplementary Text 1: Stable phase progressions dissociate oscillatory from non-oscillatory events

With the high temporal resolution of electrophysiological measures, we can dynamically assess neuronal activity while it unfolds. A growing body of literature has shown that transient high-amplitude events interfuse sustained bandlimited neuronal activity and could aid long-distance communication[1–4]. However, not every increase in bandlimited power is automatically an oscillatory event[2,5,6] because neuronal activity also features meaningful and time-varying aperiodic activity[6–9].

We outline the dissociation between oscillatory and aperiodic events in the following schematic (Supplementary Fig. S1). We generated 5 seconds of random data (x, $n_{repetition}$ = 100,000; Supplementary Fig. S1a) with a pink spectrum (1/f distribution). The five-second data traces were split into 1-second windows. Window II (1–2 seconds) and window IV (3–4 seconds) were modulated in broadband amplitude (3*x) and by adding a 10 Hz sinusoidal signal (0.8*x + 0.8*sine), respectively. The modulation parameters were chosen to i) approximately match the broadband power between the random data segments and the sine modulated window (Supplementary Fig. S1b) as well as ii) approximately match bandlimited power at 8–12 Hz for both modulated windows (Supplementary Fig. S1d).

Importantly, both modulations lead to increases in bandlimited signal power (8–12 Hz, Supplementary Fig. S1c,d) and the detection of high-amplitude events (Supplementary Fig. S1e,f). We know that the high-amplitude events in one window were aperiodic (window II) and the other oscillatory (window IV). However, in empirical neuronal data we don’t have ground truth knowledge and need to make a decision about each high-amplitude event if it is more likely to be oscillatory or not.

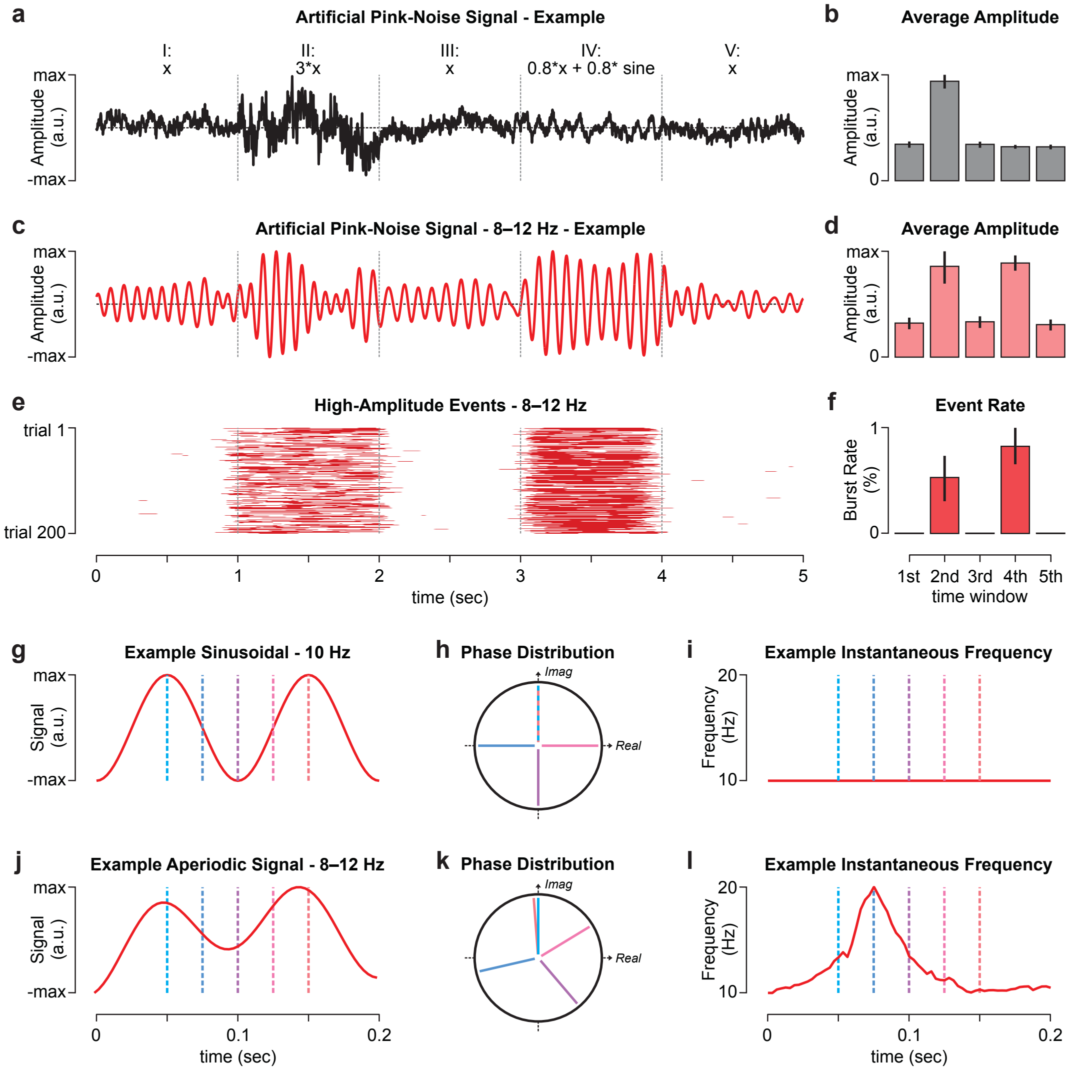


**Supplementary Figure S1 | Schematic dissociating oscillatory and aperiodic high-amplitude events. a** Example trace of a randomly generated signal with a pink (1/f) spectrum (x). The five second traces are split into five 1-second windows. Window II (1–2 seconds) and window IV (3–4 seconds) were modulated in broadband amplitude (3*x) and by adding a 10 Hz sinusoidal (0.8*x + 0.8*sine), respectively. **b** The average broadband signal amplitude strength for the middle 0.5 seconds within each window. The bars and vertical lines display the median and 25%-75% interquartile range over simulations ($n_{simulation}$ = 100,000). **c-d** Same as **a-b** for bandlimited signals filtered between 8–12 Hz. **e** The occurrence of high-amplitude events in the 8–12 Hz range (see Methods) for a subset of 200 randomly drawn trials from the simulated data traces. Red and white indicate the occurrence and absence of an event, respectively. **f** The average high-amplitude event rate for the middle 0.5 seconds within each window. **g-i** Deriving the instantaneous frequency **i** from a sinusoidal oscillatory wave form **g** using the derivative of the phase distribution **h** in the complex plane. **j-l** Same as **g-i** for an example trace of a bandlimited (8–12 Hz) signal filtered from a broadband signal with an aperiodic spectrum (1/f).

There are two caveats that hinder dissociating oscillatory and aperiodic events empirically. First, events are short, in the order of 10s to a few 100s of milliseconds[1,10]. This is a problem because in theory estimating the full power-spectrum could dissociate oscillatory (frequency-limited peak) and aperiodic events (broadband offset). Yet, to estimate a sufficient spectrum we need to include data several times the average event lengths to make predictions about bandlimited power differences thus diluting oscillatory power increases. Second, the oscillation frequency is unknown and potentially variable between events[11]. This impedes time-window-based analyses that would need to be matched to the oscillatory frequency[3,12,13].

We propose estimating the instantaneous frequency to address these two problems and determine if a given event is more likely oscillatory or aperiodic (Supplementary Fig. S1g-i). The frequency of a signal is the speed with which the phase progresses through the oscillatory cycle. When a cycle takes long the frequency is low. Similarly, the instantaneous frequency (IF) can be estimated from the temporal derivative of the signal phase: When a phase step is long between time points, IF is high. For example, in a pure sinusoidal 10 Hz wave (Supplementary Fig. S1g-i) one full oscillation takes 0.1 seconds. The phase hereby evenly progresses over the cycle, i.e., the phase angle between neighboring timepoints is constant and in the same way IF is constant at 10 Hz. Now, for a bandpass filtered aperiodic signal on the other hand, we will observe variability in the phase progression speed, i.e., the instantaneous frequency (Supplementary Fig. S1j-l).

Thus, for any high-amplitude event we can define if it is more likely oscillatory or aperiodic by quantifying the stability of the instantaneous frequency (compare Supplementary Fig. S1i versus S1I). In principle, we only need two data samples to estimate IF, which makes it suitable for very short event durations. Moreover, the approach is robust to variable frequencies between events because we assessed the IF stability, i.e., frequency difference over time, and not the exact frequency.

Overall, the schematic shows that both aperiodic and oscillatory power increases can lead to the detection of high-amplitude events in bandlimited neuronal signal. Characterizing every high-amplitude event as an “oscillatory event” is misleading. We propose IF stability as a useful measure to classify high-amplitude events as likely oscillatory or aperiodic. We used this approach to show how oscillatory high-amplitude events can have distinct effects on various measures of local neuronal activity (Supplementary Fig. S2), coupling between distant cortical areas (Supplementary Fig. S6) and behavior (Fig. 2-4).

#### Supplementary Text 2: Local high-amplitude oscillatory event-rate peaks between 8–13 Hz

We applied oscillatory and aperiodic high-amplitude event detection using instantaneous frequency stability on source-reconstructed cortical activity over a broad range of frequencies (2–64 Hz, including the delta, theta, alpha, beta and gamma ranges). To validate our detection approach, we assessed the distributions of local high-amplitude oscillatory events. We found that the mean oscillatory event rate was highest in the alpha and low beta frequency range (8–16 Hz, Supplementary Fig. S2a). Oscillatory- were also more prominent than aperiodic events for all frequencies below the gamma range (Supplementary Fig. S2b) and more likely than expected from random fluctuations ($z_{vs.random} > 6$; Supplementary Fig. S3).

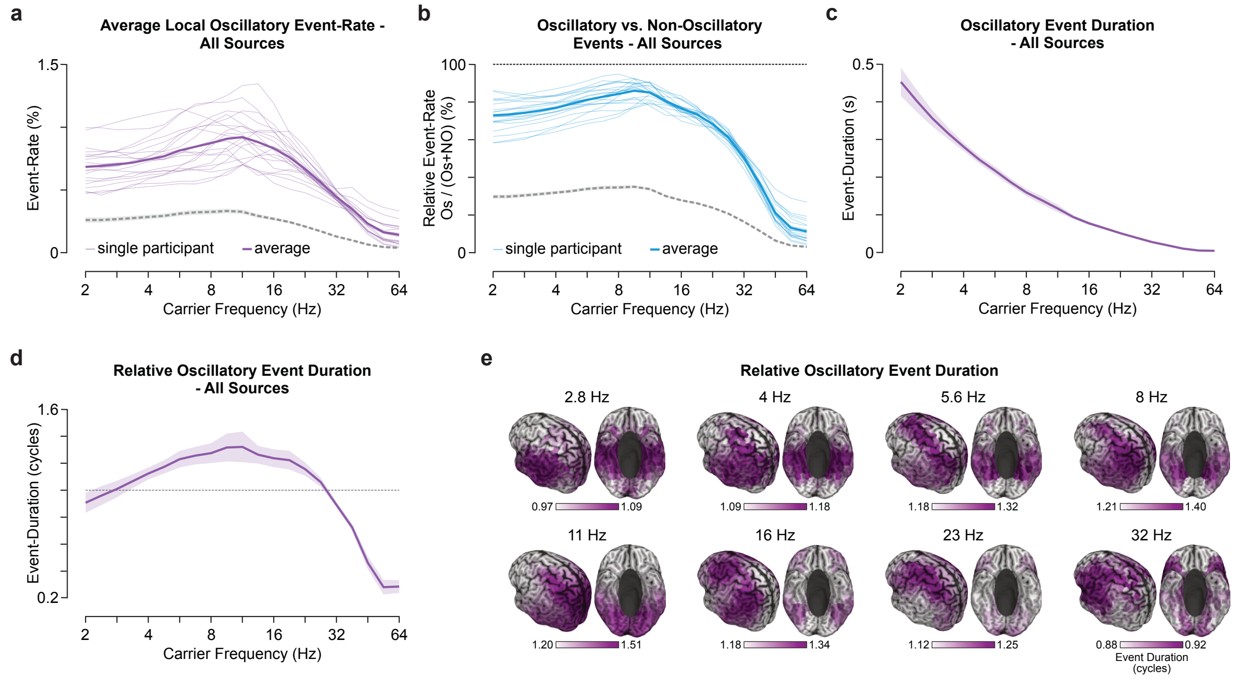


**Supplementary Figure S2 | Local high-amplitude oscillatory event-rate during task. a** Spectrum of the local oscillatory event-rate in percent averaged over time and cortical sources. Thick and thin lines indicate the group average and single participants ($n_{participant} = 20$), respectively. **b** Spectrum of the percent local oscillatory- with respect to all high-amplitude events averaged over time and space. **c** Spectrum of the event duration in seconds averaged over all participants. Thick lines with shaded areas indicate the mean and standard deviation over cortical sources. **d** Spectrum of the relative event length in cycles of the underlying carrier frequency averaged over all participants. **e** Cortical distribution of the relative event length for frequencies between 2.8–32 Hz. The color-scale indicates the $20^{th}$ to $95^{th}$ percentile range within each carrier frequency.

The median event duration was inversely related to the carrier frequency (Supplementary Fig. S2c). This observation is a consequence of methodological considerations, i.e., adaptive filter width, and intrinsic timescales of neuronal activity fluctuations[7]. We thus further quantified the relative duration (in cycles) and found that in the theta to beta frequency range the median event lasted longer than one cycle (Supplementary Fig. S2d). The relative durations for different cortical areas further highlighted frequency-specific distributions (Supplementary Fig. S2e). For example, in the high alpha range (11 Hz) relative durations peaked in the visual cortex and parietal areas. Whereas, in the theta (5.6 Hz), lower alpha (8 Hz) and particularly beta frequency range (16–23 Hz) oscillatory event activity lasted the longest in a fronto-parietal network.

Further, comparing the results against local high-amplitude events without IF stability, i.e., aperiodic events, we observed marked differences (Supplementary Fig. S4). First, aperiodic event-rates increase with higher frequencies and displays below-expected rates from the delta up to the beta frequency range. Second, the duration of individual events is considerably shorter for these lower frequencies as well (compare Supplementary Fig. S2c,d and Supplementary Fig. S4c,d). Third, the cortical distribution of relative event durations displays little spectral specificity up to 32 Hz and was not or anti-correlated with oscillatory event durations in the delta, theta, and alpha frequency range.

The above description of local high-amplitude oscillatory events is a first step to validate our detection approach. For example, we observed that oscillatory events were strongest and most persistent in the alpha frequency range and further reproduced high oscillatory event-rates in the beta range[1,3,13]. Generally, local high-amplitude oscillatory events appear to be a sensible measure for neuronal activity overlapping with previous findings on cortical oscillatory generators[14–16].

**Supplementary Text 3: Time-averaged oscillatory event coincidences predict amplitude coupling**

One question within the current study was to determine if coincident high-amplitude oscillatory events (Supplementary Fig. S5) could shape amplitude coupling networks[15,17,18]. However, we first assessed whether oscillatory event coincidences (OEC) are a sensible and reliable measure of functional coupling. We generated cortical OEC patterns for three regions of interest in the (left) auditory, somatosensory and medial prefrontal cortex at 11 Hz (Supplementary Fig. S6a; one-sided t-test, df = 19, $p_{FDR} < 0.05$). The patterns display sensible[17] coupling with neighboring sources but also long-distance coupling to homologue areas in the other hemisphere (ISSC) and intra-hemispherically between medial prefrontal and lateral parietal areas. The full connectome displayed strong intra- but also inter-hemispheric connections with OEC exceeding random co-occurrences by a factor of 2.5–6.5 (Supplementary Fig. S6b) and amplitude-coupling matched surrogate data by a factor of 1.2–1.9 (Supplementary Fig. S7). The patterns were further reliable between participants at a level of $r_{11Hz,reliability} = 0.40$ (reliability: $range_{2\text{-}45Hz} = 0.20–0.42$; Supplementary Fig. S6c). Generally, we found the strongest global coupling, i.e., degree, for lateral parietal and superior temporal areas between 4–23 Hz with spectrally evolving spatial peaks from more ventral to dorsal areas with increasing frequency. In higher frequencies the spatial distribution overlapped with hubs of residual muscle activity[19].

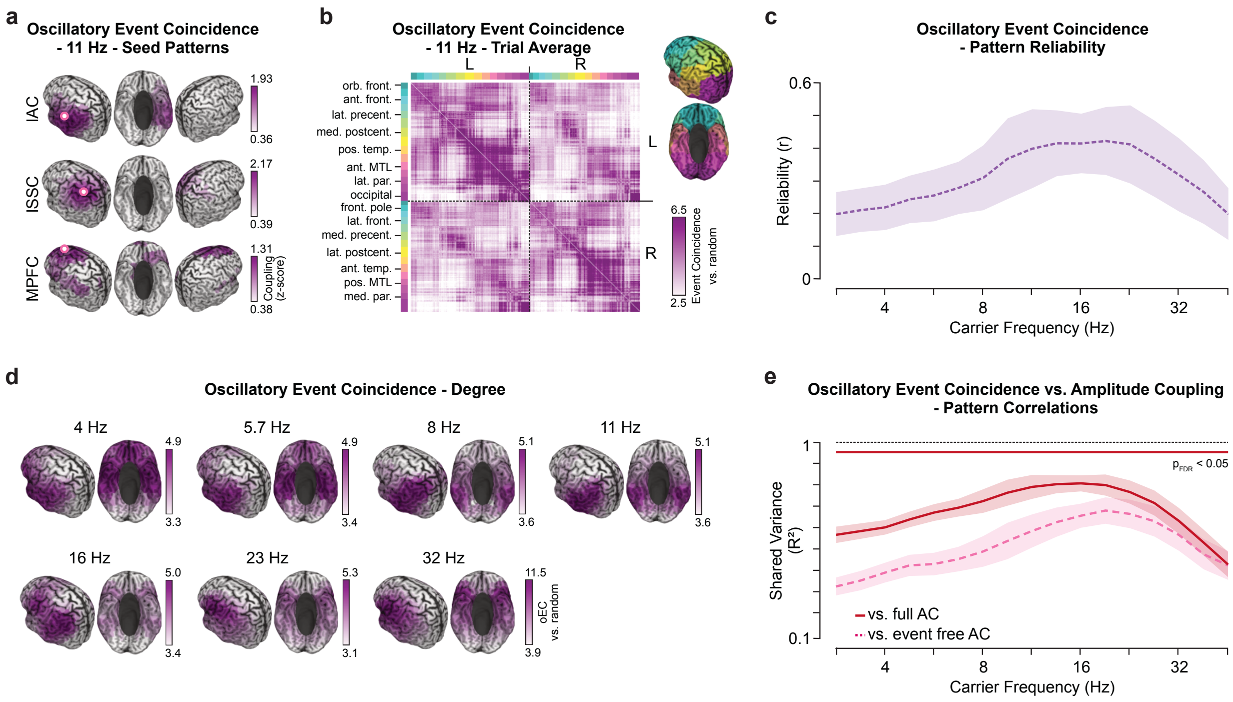


**Supplementary Figure S6 | Time-averaged oscillatory event coincidences and the relationship to amplitude coupling. a** Oscillatory event coincidence (OEC) patterns for seeds in the left auditory cortex (lAC, [-54, -22, 10]), left somatosensory cortex (lSSC, [-42, -26, 54]), medial prefrontal cortex (MPFC, [-3, 39, -2])[17] at 11 Hz. Coupling values were z-scored over the cortex within each participant ($n_{participant}$ = 20) and t-tested against zero (one-sided, unpaired, FDR-corrected at $p_{FDR}$<0.05). Non-significant values were omitted. **b** Full adjacency matrix for all cortical connections at 11 Hz. The connection strength is indicated as coincidence rate against random (random reassignment of trials between connected sources). Capital letters L and R denote sources within the left and right hemisphere, respectively. Sources were color-coded according to anatomical regions, depicted as insert. **c** Spectrum of the reliability of cortical OECs between participants. Thick lines with shaded areas indicate the mean and standard deviation over the cortex, respectively. **d** Degree, i.e., the average OEC versus random of each source, at frequencies between 4–32 Hz. All color-scales were set between the 5th and the 95th percentile. **e** Spectrum of the shared variance, i.e., the squared correlation coefficient, for the relationship between OECs and amplitude coupling within participants. The correlations were computed while including (solid line) or excluding (dashed line) high-amplitude event time points. Thick lines with shaded areas indicate the mean and standard deviation over the cortex, respectively. The thick bar on top denotes a significant difference between shared variances (Wilcoxon signed rank test on cortical average, df=19, $p_{FDR}$<0.05).

Now, are the observed networks compatible with amplitude coupling[15,17,18] (Supplementary Fig. S8)? As a first piece of evidence, we directly correlated networks defined by time-averaged OECs and time-averaged amplitude coupling, respectively, within every participant and frequency (Supplementary Fig. S6e, solid line). We identified high correlations between both coupling measures ranging from 50–90% shared variance (correlation squared) for frequencies up to 30 Hz.

Amplitude coupling networks, when averaged over time, also include time points of high-amplitude oscillatory and aperiodic events interleaved with low-amplitude activity. We thus computed event free, i.e., sustained, amplitude coupling. If OECs can describe distinct network states, the relationship to amplitude coupling would qualitatively change. If, on the other hand, high-amplitude oscillatory events only have a quantitative effect we would expect a lower amplitude coupling magnitude but the same spatial network topography. We found evidence for the former, distinct network states associated with high-amplitude oscillatory transients: The similarity between event free amplitude coupling (Supplementary Fig. S6e, dashed line) and OEC patterns decreased over the entire spectrum (Wilcoxon signed rank test(19), $p_{FDR} < 0.05$). Further, the lower similarity cannot be explained by a drop in reliability (Supplementary Fig. S9).

We further scrutinized the relationship between transient aperiodic event coincidence and amplitude coupling (Supplementary Fig. S10). The results displayed that (i) aperiodic event coincidence patterns appeared to be more focal, (ii) the degree peaks shifted to frontal regions (theta to alpha frequencies), and (iii) the similarity with amplitude coupling networks was decreased for low (theta and alpha, 4.8–13 Hz) but increased for high frequencies (27–45 Hz, paired t-test(19) on connectome averaged similarity, $p_{FDR} < 0.05$).

Conclusively, coinciding oscillatory high-amplitude transients revealed a reliable functional network organization. The identified networks strongly overlapped with time-averaged amplitude coupling in the delta to high beta frequency ranges: Between 50–90% of an amplitude coupling pattern can be explained from analyzing less than 5% of the data, i.e., high-amplitude oscillatory events. Moreover, excluding high-amplitude events from amplitude coupling qualitatively altered coupling patterns, predominantly for lower frequencies. The findings indicate initial evidence for distinct network states during transient high-amplitude oscillatory events and sustained low-amplitude periods.

**Supplementary Text 4: Coincidences of high-amplitude oscillatory events are dynamically modulated during task performance**

So far, we estimated time-averaged functional coupling neglecting dynamic changes over time. But with our decision-making task we can dynamically resolve coupling changes to intrinsic and exogenous variables[20]. Each trial consisted of three distinct epochs, i.e., pre-stimulus (-1.5–-1 s), sensory- (-1–0 s) and decision-making phase (0–2.8 s with respect to the framing cue). From the pre-stimulus to the sensory-phase the visual stimulation changed, i.e., targets appeared on screen. Between the sensory- and decision-phase the visual stimulation was identical (with the exception of the 0.3 s framing cue) but the task demand and related cognitive processing changed, e.g., the exploration of targets versus decision-related information sampling[20]. Thus, we used the second half of the sensory-phase as reference, i.e., baseline period (-0.5–0 s from framing), to exploratively assess coupling changes to visual stimulation and cognition.

The results showed that local high-amplitude oscillatory event-rates as well as transient network activity (Supplementary Fig. S11 and S12) were modulated by the visual stimulus. We found relatively increased OEC during the pre-stimulus phase and decreased upon visual stimulus onset (Supplementary Fig. S12a,b). Stimulation induced decreases in neuronal activity, particularly in sensory related areas and frequencies, is an established phenomenon that has been reported repeatedly[21]. This effect was already present in local event activity (Supplementary Fig. S12a,c). Further, visual stimulation led to a transient decrease in the oscillatory/aperiodic event ratio between 0.3–0.5 seconds after both stimulus onset and framing cue onset (Supplementary Fig. S12b,d).

Brain-wide oscillatory event coincidences further displayed a systematic relationship with the decision-making task. During the decision phase we observed both increased and decreased OEC for a variety of frequencies and time points. Functional coupling relatively increased to the sensory baseline in the theta to beta frequency range from around 1 second post-framing onwards

(Supplementary Fig. S11b; paired t-test(19) against baseline, p<0.01). Decreased OEC on the other hand occurred predominantly within the first 0.7 seconds post-framing and in the beta frequency range as well as around stimulus offset in the gamma range (Supplementary Fig. S11b, right panel). We identified compatible spectro-temporal profiles for dynamic amplitude coupling while including (Supplementary Fig. S13) or excluding high-amplitude events (Supplementary Fig. S14).

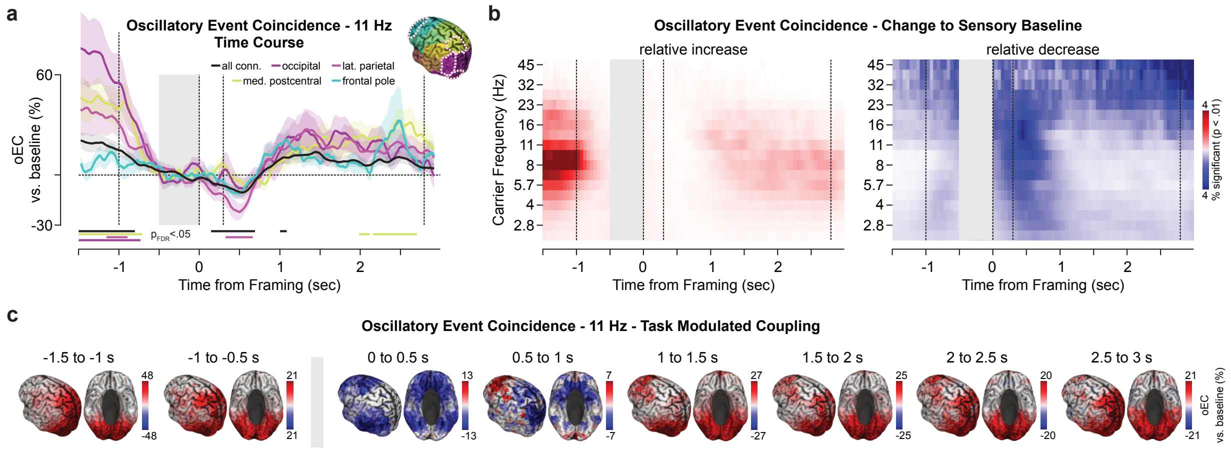


**Supplementary Figure S11 | Oscillatory event coincidence dynamics during decision-making. a** Trial dynamics of oscillatory event coincidences at 11 Hz averaged over participants (n = 20). The event coincidence rate was normalized by the average over the sensory baseline (-0.5 to 0 s from framing). Displayed are dynamic OEC rates averaged over all connections (black) and within the occipital- (dark purple), lateral parietal (purple), medial postcentral cortex (green) and the frontal pole (cyan). The vertical lines indicate the stimulus onset, framing cue onset and offset as well as the stimulus offset (left to right). The inset displays the analyzed cortical sources highlighted (in pink) on the left hemisphere. Thick lines depict significant changes with respect to the sensory baseline (two-sided t-test(19), $p_{FDR}$<0.05, FDR-corrected over time). **b** Number of significantly modulated connections with respect to the sensory baseline (grey bar) for relative increases (left) and decreases (right). Significantly modulated connections (two-sided t-test(19), uncorrected p<0.01) were computed at each time point and for frequencies between 2.3–45 Hz against the average coupling within the baseline window over participants. **c** Average change in coupling with respect to the sensory baseline for each cortical source. The respective changes were averaged within each window and over all connections from each source.

Moreover, the network of modulated OEC connections varied over time. Exemplary, we visualized OEC dynamics at 11 Hz, i.e., a frequency with consistent high-amplitude oscillatory events (see Supplementary Fig. S2). First, we identified coupling changes related to visual stimulation in a network including early visual and medial parietal areas (Supplementary Fig. S11c, -1.5 to -1 s from framing). Second, post framing decreases highlighted a network of ventral visual areas and the medial prefrontal cortex (0–0.5 s from framing). Third, OEC increased during the later stages of the decision phase (1–3 s from framing) and displayed network transitions from ventral visual and lateral prefrontal connections (1–2 s) to extrastriatal visual and medial parietal cortex connections (1.5–3 s from framing).

In sum, we found that the probability of high-amplitude oscillatory event coincidences was dynamically modulated during a demanding decision-making task. The dynamics were hereby compatible with amplitude coupling further validating the approach. Distinct neuronal processing demands between the sensory baseline and the decision phase of the trial (1-2.5s post-framing) were evident in increased OEC in the theta, alpha and beta frequency ranges. Moreover, the modulated connections displayed network transitions in line with routing internal and external information[22,23] with respect to task demands, i.e., from for example attention maintenance[24] to evidence accumulation and motor preparation[25,26].

### Supplementary Figures

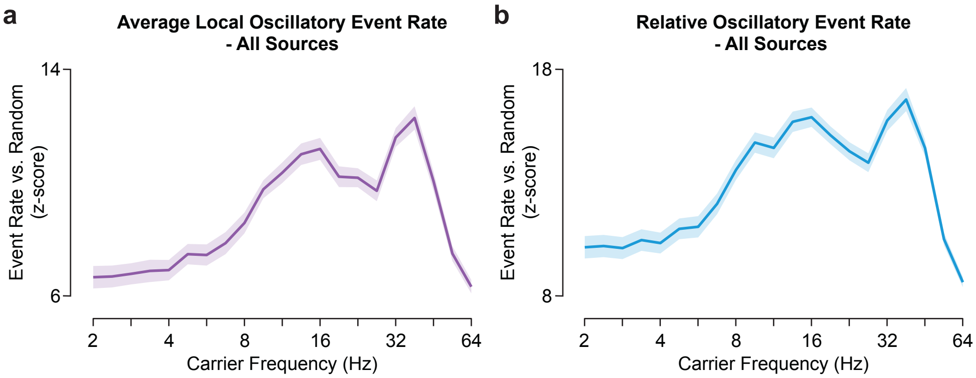


**Supplementary Figure S3 | Local oscillatory event rates against random.** Spectral distribution of local event rates normalized (z-scored) to random occurrences for **a** the absolute oscillatory event rate and **b** the oscillatory event rate relative to all high-amplitude events (oscillatory / (oscillatory + aperiodic)). Thick lines and shaded areas indicate the mean over participants (n = 20) and the SEM.

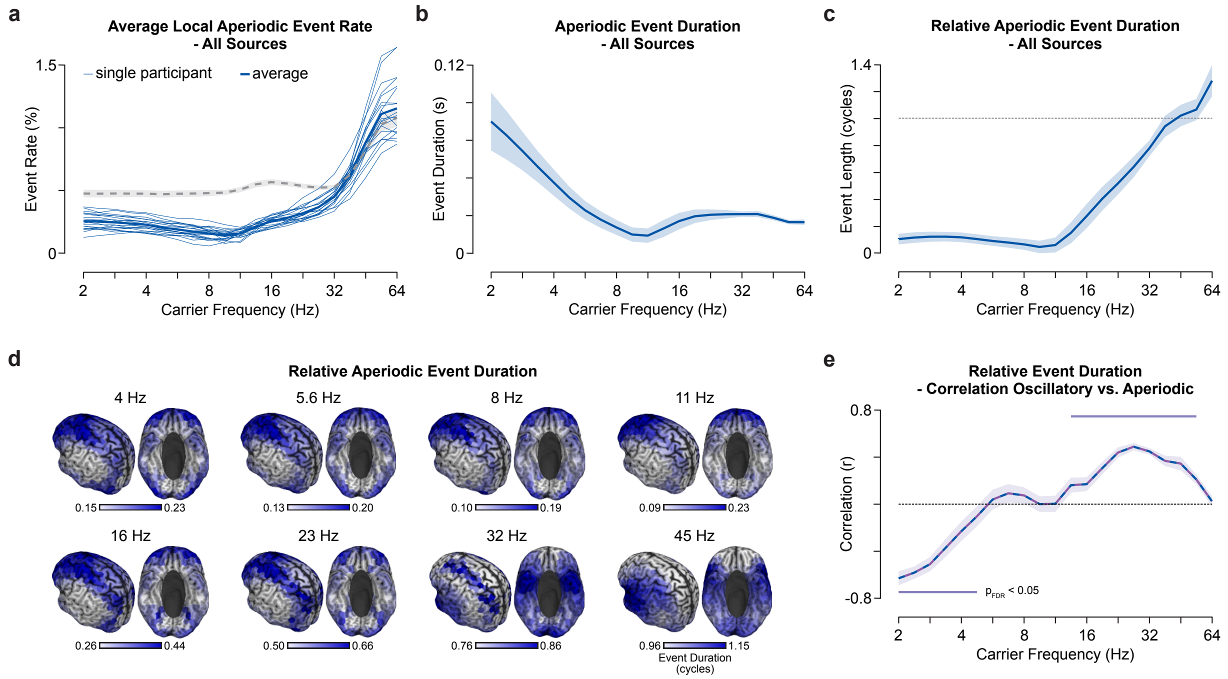


**Supplementary Figure S4 | Local aperiodic high-amplitude event rates during task. a** The spectrum of the local aperiodic burst rate in percent averaged over time and cortical sources. Thick and thin lines indicate the group average and single participants, respectively. The dashed gray line depicts event rate under random conditions. **b** Spectrum of event durations in seconds averaged over all participants. Thick lines with shaded areas indicate the mean and standard deviation over cortical sources. **c** Spectrum of the relative event duration in cycles of the underlying carrier frequency averaged over all participants. **d** Cortical distribution of the relative aperiodic event durations for frequencies between 4–45 Hz. The color-scale indicates the 20$^{th}$ to 95$^{th}$ percentile range within each carrier frequency. **e** Spectrum of the (within participant) correlation of the spatial distributions of relative event durations between oscillatory vs. aperiodic events (Supplementary Fig. S2e and Supplementary Fig. S4d). Thick lines with shaded areas indicate the mean and standard error over participants. The values were Wilcoxon rank-tested (df = 19) against zero ($p_{FDR}$<0.05).

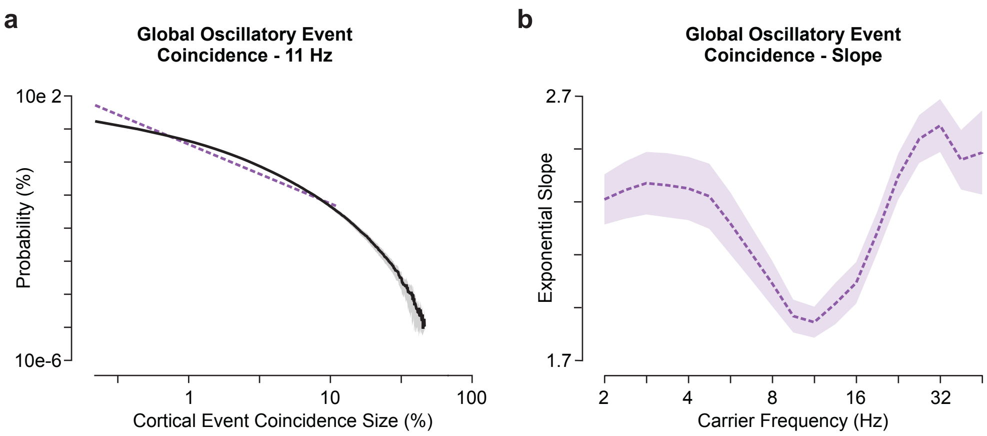


**Supplementary Figure S5 | Global oscillatory event coincidences. a** The histogram of cortical oscillatory event coincidence sizes at 11 Hz in log-log scale. The purple dashed line displays the fit of the exponential decay of cortical event coincidence sizes between 1 and 10% sources. **b** Spectrum of the slope of the fitted exponential decay of cortical oscillatory event coincidence sizes. Thick lines with shaded areas indicate the mean and standard error over participants. Global event coincidences represent an intermediate step between local events and assessing co-occurrence of events in networks. The measure describes the number of cortical sources displaying high-amplitude events locally for each moment in time and can range between no source (size = 0%) and all sources displaying events (size = 100%).

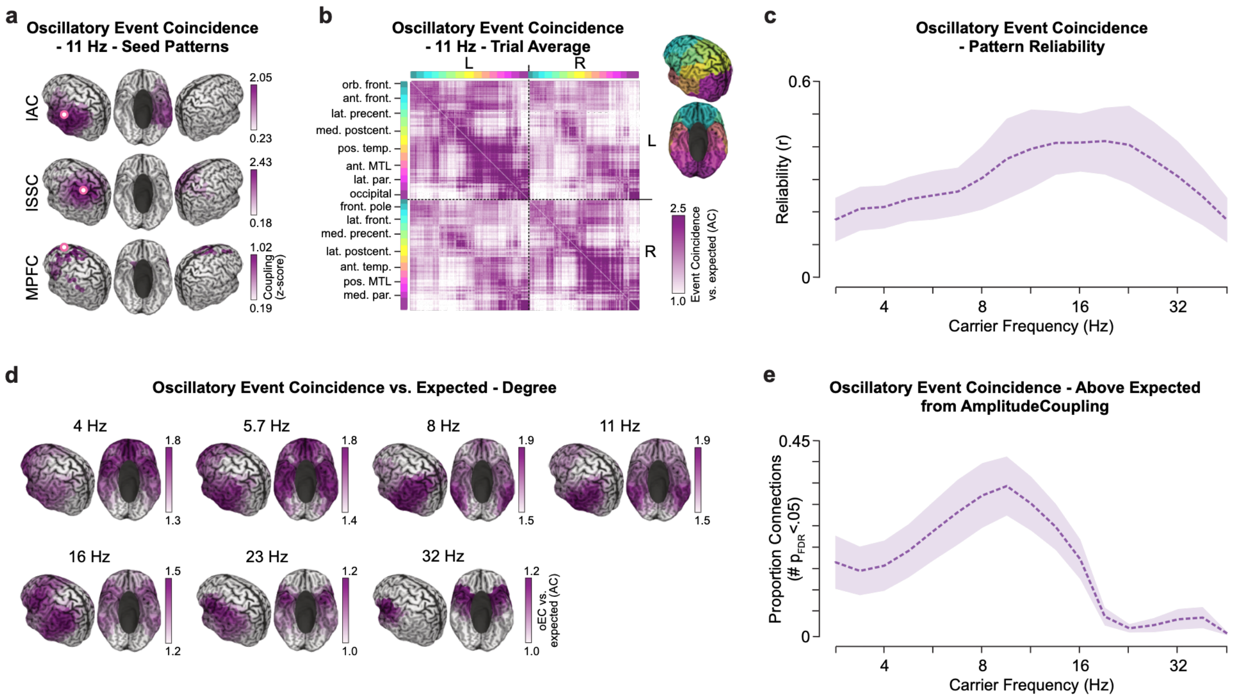


**Supplementary Figure S7 | Time-averaged oscillatory event coincidences and the relationship to amplitude coupling.** This is an extension to Supplementary Fig. S6 for oscillatory event coincidence rates tested against rates expected under matched amplitude coupling conditions. Here, for each connection, participant ($n_{participant}$ = 20) and frequency we simulated coincident event rates based on pink noise amplitude envelopes with amplitude correlation equal to the empirical value ($n_{repetition}$ = 5000). **a** Oscillatory event coincidence (OEC) patterns for seeds in the left auditory cortex (lAC, [-54, -22, 10]), left somatosensory cortex (lSSC, [-42, -26, 54]), medial prefrontal cortex (MPFC, [-3, 39, -2])[17] at 11 Hz. Coupling values were z-scored over the cortex within each participant and t-tested against zero (one-sided, unpaired, FDR-corrected at $p_{FDR}<0.05$). Non-significant values were omitted. **b** Full adjacency matrix for all cortical connections at 11 Hz. The connection strength is indicated as coincidence rate against expected from amplitude coupling (1 = expected value). Capital letters L and R denote sources within the left and right hemisphere, respectively. Sources were color-coded according to anatomical regions, depicted as insert. **c** Spectrum of the reliability of cortical OECs between participants. Thick lines with shaded areas indicate the mean and standard deviation over the cortex, respectively. **d** Degree, i.e., the average OEC versus expected of each source, at frequencies between 4–32 Hz. All color-scales were set between the 5$^{th}$ and the 95$^{th}$ percentile. **e** Spectrum of the number of connections with coincidence event rates above expected (one-sided t-test(19), $p_{FDR} < 0.05$). Dashed line with shaded area indicates the mean and standard deviation over the cortex, respectively.

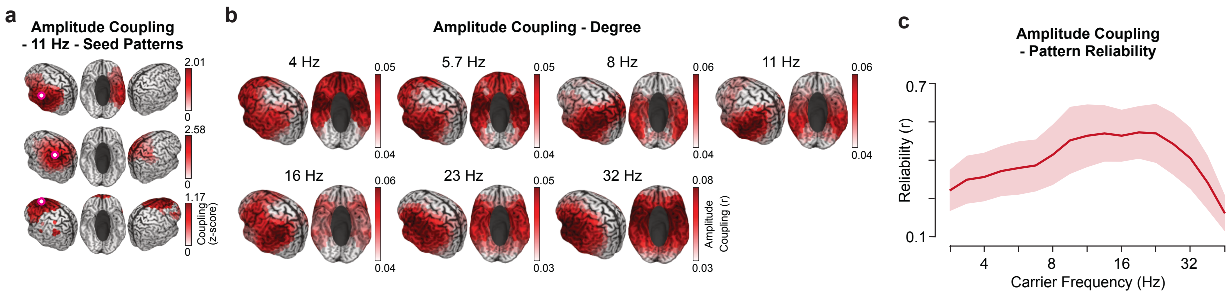


**Supplementary Figure S8 | Time-averaged amplitude coupling.** The Figure extends the findings from Supplementary Fig. S6 to full amplitude coupling (including high-amplitude events). The applied methodology is compatible. **a** Amplitude coupling patterns for seeds in the left auditory cortex (lAC, [-54, -22, 10]), left somatosensory cortex (lSSC, [-42, -26, 54]) and medial prefrontal cortex (MPFC, [-3, 39, -2]) at 11 Hz. Coupling values were z-scored over the cortex within each participant and t-tested against zero (one-sided, unpaired, FDR-corrected at $p_{FDR}$<0.05). Non-significant values were omitted. **b** The degree, i.e. the average coupling strength of each source, at frequencies between 4–32 Hz. All color-scales were set between the 5$^{th}$ and the 95$^{th}$ percentile within each panel. **c** Spectrum of the reliability of amplitude coupling patterns between participants. Thick lines with shaded areas indicate the mean and standard deviation over the cortex, respectively.

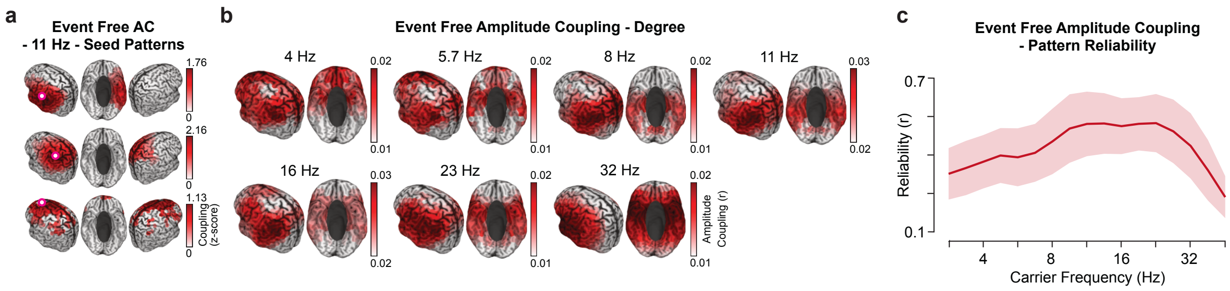


**Supplementary Figure S9 | Time-averaged event free amplitude coupling.** The Figure extends the findings from Supplementary Fig. S6 to sustained, i.e. event free, amplitude coupling (AC). The applied methodology is compatible. **a** Amplitude coupling patterns for seeds in the left auditory cortex, left somatosensory cortex, medial prefrontal cortex at 11 Hz. Coupling values were z-scored over the cortex within each participant and t-tested against zero (one-sided, unpaired, FDR-corrected at $p_{FDR}<0.05$). Non-significant values were omitted. **b** The degree, i.e. the average coupling strength of each source, at frequencies between 4–32 Hz. All color-scales were set between the 5$^{th}$ and the 95$^{th}$ percentile within each panel. **c** Spectrum of the reliability of amplitude coupling patterns between participants. Thick lines with shaded areas indicate the mean and standard deviation over the cortex, respectively.

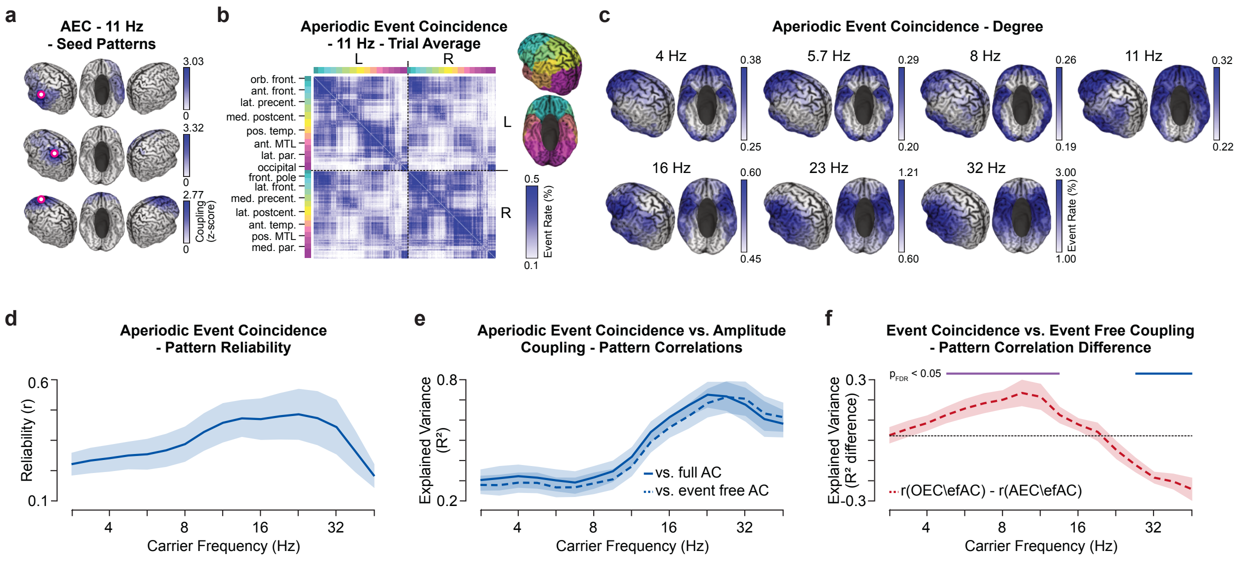


**Supplementary Figure S10 | Time-averaged aperiodic event coincidences and comparison to other coupling measures.** The Figure extends the findings from Supplementary Fig. S6 to aperiodic event coincidences (AEC). The applied methodology is compatible. **a** AEC patterns for seeds in the left auditory cortex, left somatosensory cortex, medial prefrontal cortex at 11 Hz. Coupling values were z-scored over the cortex within each participant and t-tested against zero (one-sided, unpaired, FDR-corrected at $p_{FDR}<0.05$). Non-significant values were omitted. **b** The full adjacency matrix for all cortical connections at 11 Hz. Capital letters L and R denote sources within the left and right hemisphere, respectively. Sources were color-coded according to anatomical regions, depicted as insert. **c** The degree, i.e. the average AEC rate of each source, at frequencies between 4–32 Hz. All color-scales were set between the 5$^{th}$ and the 95$^{th}$ percentile within each panel. **d** Spectrum of the reliability of AEC patterns between participants. Thick lines with shaded areas indicate the mean and standard deviation over the cortex, respectively. **e** The spectrum of the shared variance, i.e. the squared correlation coefficient, for the relationship between AEC and amplitude coupling patterns within participants. Amplitude correlations were computed with (solid line) and without (dashed line, efAC) high-amplitude event time points. Thick lines with shaded areas indicate the mean and standard deviation over the cortex, respectively. **f** The spectrum of the difference in how much variance oscillatory- (Supplementary Fig. S6e, dashed line) and aperiodic event coincidence patterns (Supplementary Fig. S10e, dashed line) can explain of sustained amplitude coupling patterns ($R^2_{OEC\backslash efAC}$-$R^2_{AEC\backslash efAC}$).

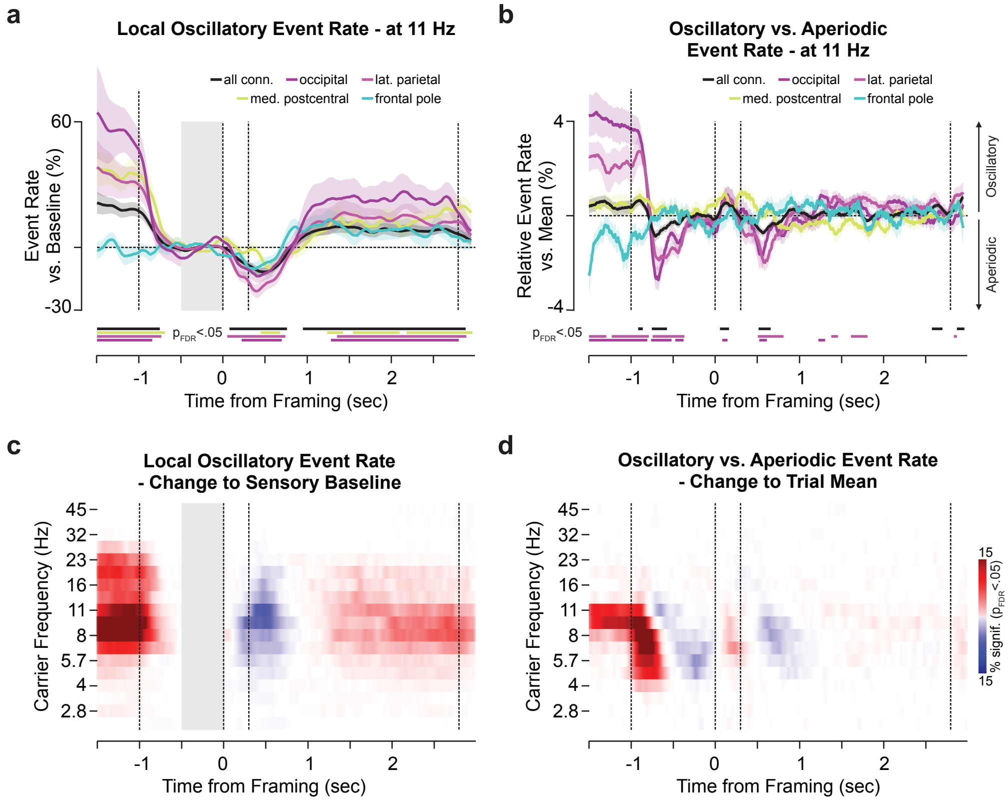


**Supplementary Figure S12 | Local event rate dynamics during decision-making. a** Local oscillatory event rates dynamics at 11 Hz. Event rates were normalized by the average over the sensory baseline (-0.5 to 0 s from framing). Displayed is the local oscillatory event rate averaged over all sources (black) and within the occipital- (dark purple), lateral parietal (purple), medial postcentral cortex (green) and the frontal pole (cyan). The vertical lines indicate the stimulus onset, framing cue onset and offset as well as the stimulus offset (left to right). Thick lines depict significant changes with respect to the sensory baseline (two-sided, unpaired t-test(19), FDR-corrected over time at $p_{FDR}<0.05$). **b** Local relative event rate (oscillatory / (oscillatory + aperiodic)) trial dynamics at 11 Hz for all sources (black) and within the same regions of interest as above. Significance (thick lines, $p_{FDR}<0.05$) was tested against the mean relative event rate during stimulus presentation (two-sided, unpaired t-test, df = 19). **c** The number of sources that displayed significantly modulated oscillatory event rates with respect to the sensory baseline (grey bar), at each time point and for frequencies between 2.3–45 Hz (t-test, df = 19, uncorrected p<0.01). **d** Number of significantly modulated relative event rates against the mean over the stimulus presentation period (-1.5 to 2.8 s from framing) for frequencies between 2.3–45 Hz.

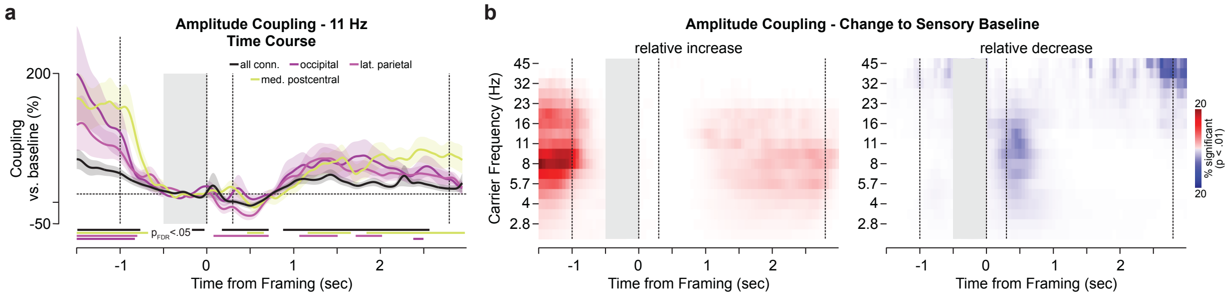


**Supplementary Figure S13 | Amplitude coupling dynamics during decision-making.** This Figure is an extension to the findings presented in Supplementary Fig. S11 for full amplitude coupling (including events). **a** Amplitude coupling trial dynamics at 11 Hz. Coupling was normalized with the average over the sensory baseline (-0.5 to 0 s from framing). Displayed is the dynamic coupling averaged over all connections (black) and within the occipital- (dark purple), lateral parietal (purple), and the medial postcentral cortex (green). The vertical lines indicate the stimulus onset, framing cue onset and offset as well as the stimulus offset (left to right). Thick lines depict significant changes with respect to the sensory baseline ($p_{FDR}<0.05$). **b** Number of significantly modulated connections with respect to the sensory baseline (grey bar) for relative increases (left) and decreases (right). Significantly modulated connections (t-test, df = 19, uncorrected p<0.01) were computed at each time point and for frequencies between 2.3–45 Hz against the average coupling within the baseline window over participants.

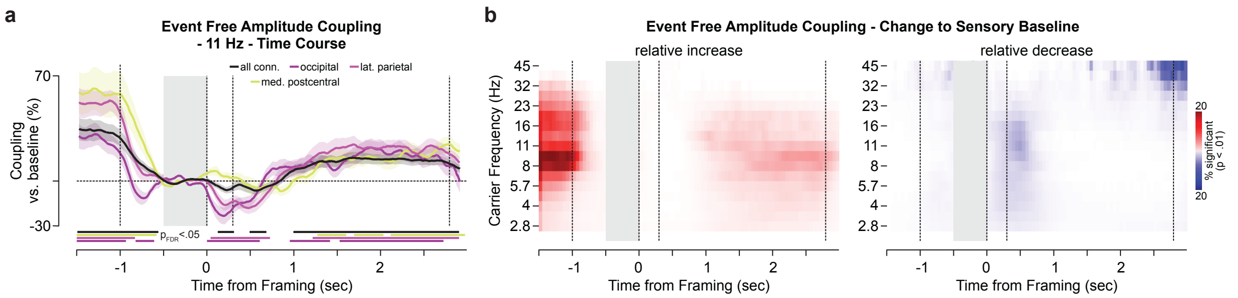


**Supplementary Figure S14 | Event free amplitude coupling dynamics during decision-making.** This Figure is an extension to the findings presented in Supplementary Fig. S11 for sustained, i.e. event free, amplitude coupling (efAC). **a** efAC trial dynamics at 11 Hz. Coupling was normalized by the average over the sensory baseline (-0.5 to 0 s from framing). Displayed is the dynamic coupling averaged over all connections (black) and within the regions-of-interest outlined above. Thick lines depict significant changes with respect to the sensory baseline ($p_{FDR}<0.05$). **b** Number of significantly modulated connections with respect to the sensory baseline (grey bar) for relative increases (left) and decreases (right). Significantly modulated connections (t-test, df = 19, uncorrected p<0.01) were computed at each time point and for frequencies between 2.3–45 Hz against the average coupling within the baseline window over participants.

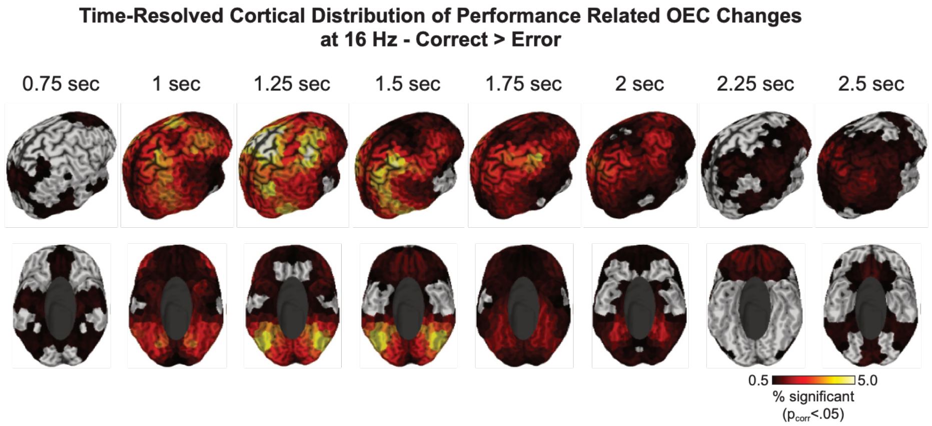


**Supplementary Figure S15 | Time-resolved cortical distribution of performance related coupling (correct > error) at 16 Hz.** Cortical distribution of the number of performance related connections per source averaged over successive time windows (window length = 0.25 s). Timing labels denote each window's starting time with respect to the framing cue onset. The color-scale displays the number of significantly increased (correct > error, two-sided paired t-test(19), cluster permutation corrected p<.05) OEC connections per time window. The results were averaged over hemispheres.

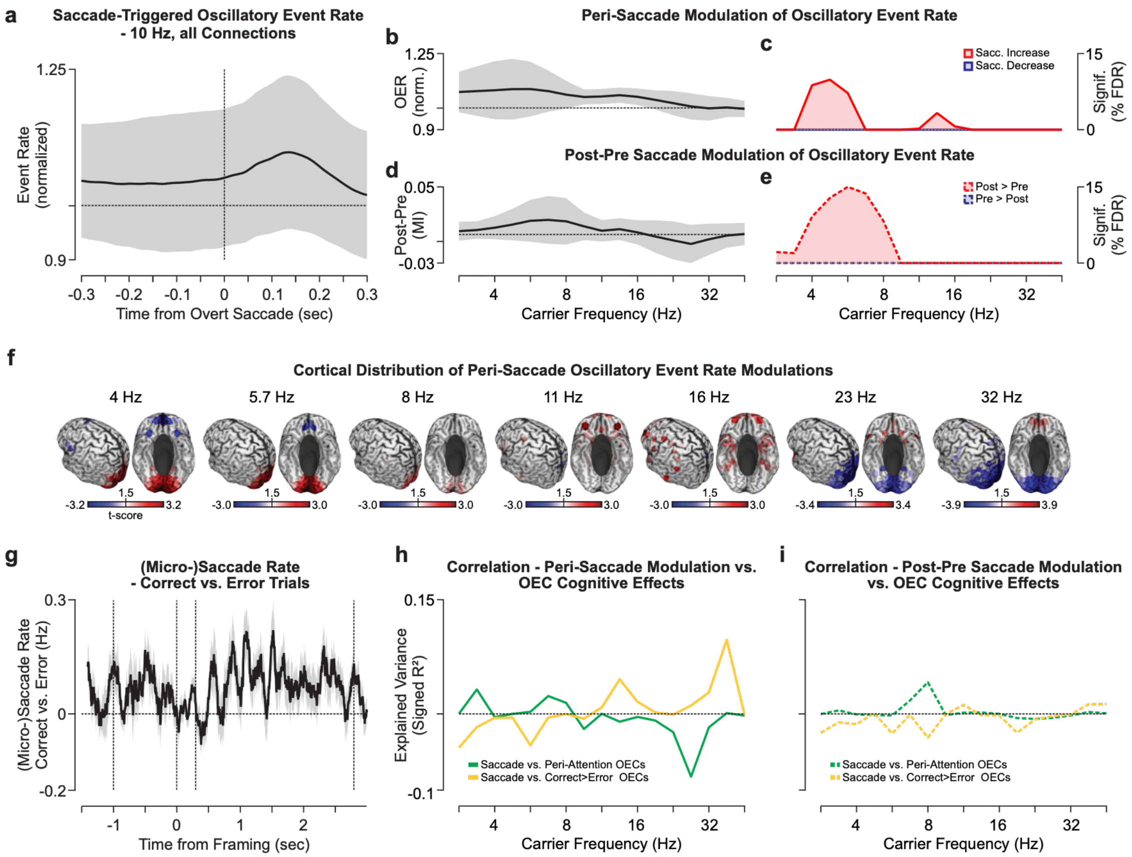


**Supplementary Figure S16 | Oculomotor related oscillatory event dynamics. a** Saccade-triggered local oscillatory event rate at 10 Hz. Displayed is the average (black line) and standard deviation (shaded area) over all sources and identified oculomotor activities (saccades and microsaccades). The data was normalized to the average event rate within each cortical source. **b-c** Peri-saccade (±0.05 s) modulation of oscillatory event rates. Displayed are the spectrum of **b** average normalized peri-saccade event rates and **c** the number of significantly modulated sources (two-sided t-test(19), p<.05, FDR-corrected). **d-e** Same as **b-c** for the post-saccadic modulation (modulation index = post-pre/(post+pre), post = 0–0.15 s, pre = -0..15–0 s). **f** Cortical distribution of peri-saccade modulated sources for frequencies between 4–32 Hz (t-scores, two-sided t-test(19), p<.05, uncorrected). Red and blue areas represent increased and decreased oscillatory event rates, respectively. **g** Performance modulated (correct vs. error) oculomotor activity over the full trial period. Thick line and shaded area denote the mean and standard error over participants ($n_{participant}$ = 20). Statistical testing (two-sided paired t-test(19)) did not show a significant difference after multiple comparison correction (FDR-correction). **h-i** Correlating the spatial distribution of **h** peri-saccade and **i** post-saccadic event rate changes with peri-attention (green line, see Fig. 4) and performance related (orange line, see Fig. 3) modulations of oscillatory event coincidences for frequencies between 2.8–45 Hz. Plotted is the signed squared correlation coefficient.

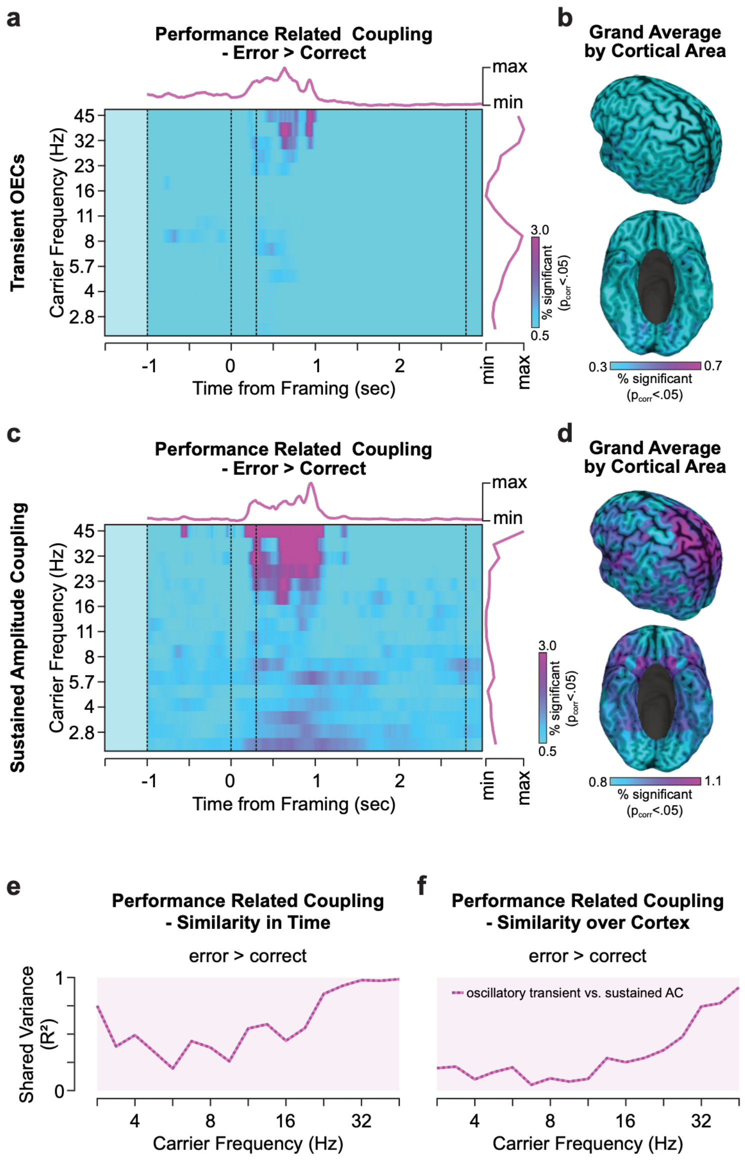

**Supplementary Figure S17 | Spectral and spatial dissociation of error-related amplitude coupling networks.** This Figure is an extension to the findings presented in Fig. 3 for coupling increased during error trials (error > correct). **a** Time and frequency resolved distribution of performance related (error > correct, one-sided t-test(19), cluster permutation corrected p<.025) transient oscillatory coupling connections. Colored lines on top and to the right denote the average over frequencies and time, respectively. **b** Average over the full time and frequency resolved distribution of behaviorally relevant transient oscillatory coupling modulations within each cortical source. **c-d** Same as **a-b** for sustained (event-free) amplitude coupling. **e-f** Pearson correlation of the behavioral coupling effects between transient oscillatory and sustained amplitude coupling comparing the **e** temporal (see panel a and c) and **f** spatial (Supplementary Fig. S19) distribution of effects. The values depict the shared variance ($R^2$) within each frequency.

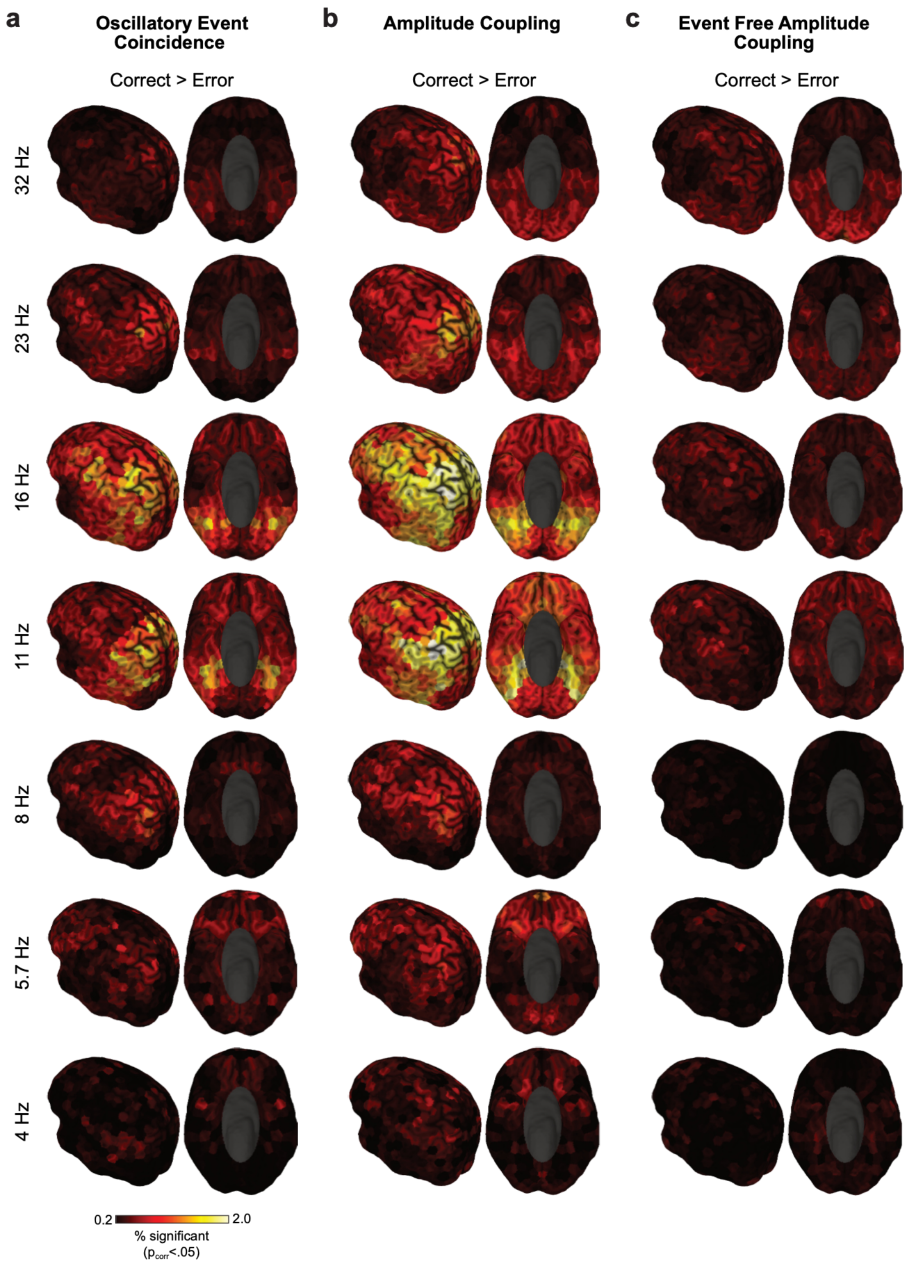


**Supplementary Figure S18 | Cortical distribution of performance related coupling (correct > error).** The number of functional coupling connections significantly increased during correct vs. error trials averaged over connections and time points per source and carrier frequency (4–32 Hz, bottom to top, correct > error, paired t-test, df = 19, cluster permutation corrected p<0.05). Separately for **a** oscillatory event coincidences, **b** amplitude coupling and **c** sustained (event free) amplitude coupling. The color scale was set from 0.2% to 2.0% significant connections and time points, according to the 5$^{th}$ and 95$^{th}$ percentile over all frequencies for oscillatory event coincidence effects (correct > error, cluster permutation corrected p<0.05).

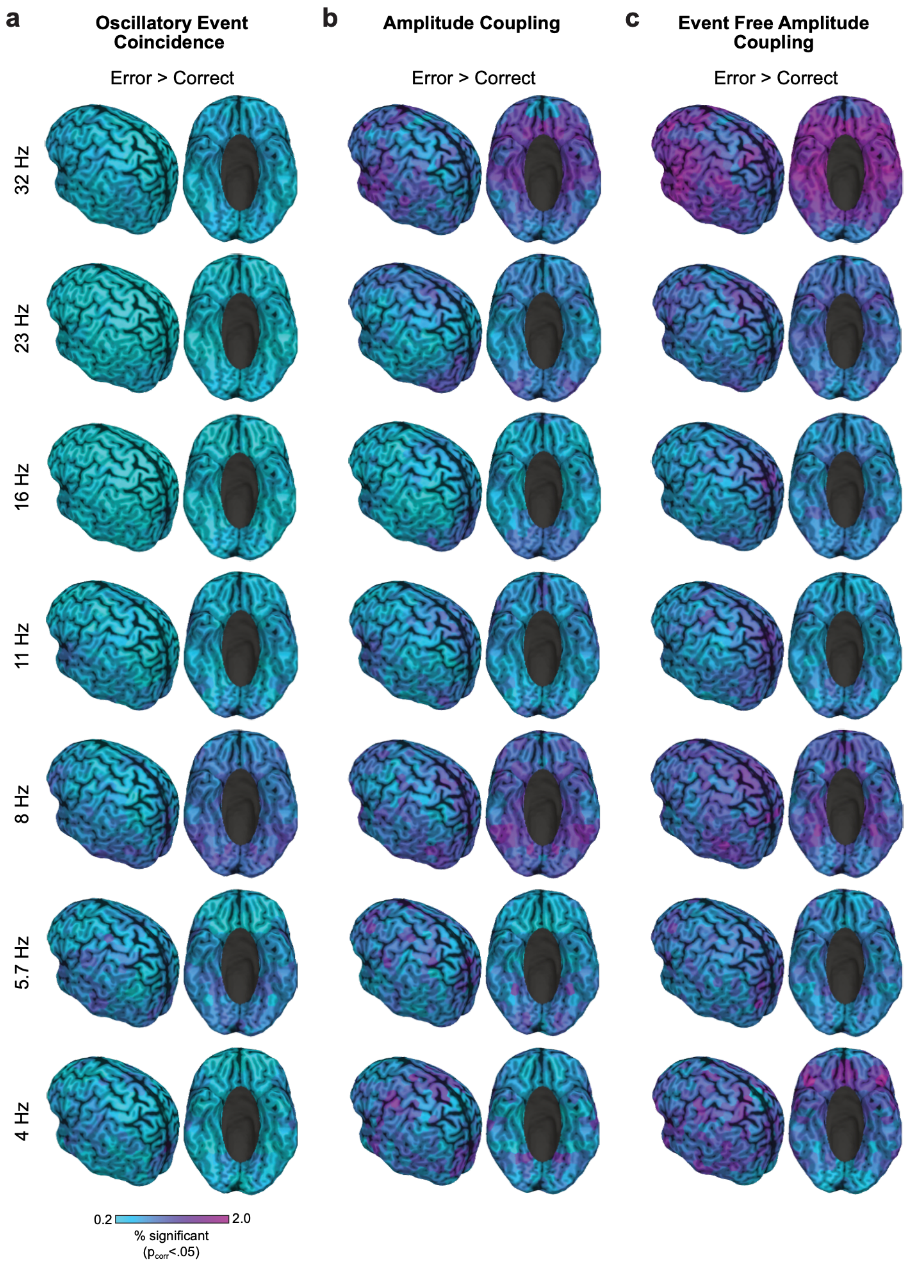


**Supplementary Figure S19 | Cortical distribution of performance related coupling (error > correct).** The number of functional coupling connections significantly increased during correct vs. error trials averaged over connections and time points per source and carrier frequency (4–32 Hz, bottom to top, error > correct, paired t-test, df = 19, p<0.05). Separately for **a** oscillatory event coincidences, **b** amplitude coupling and **c** event free amplitude coupling. The color scale was set from 0.2% to 2.0% significant connections and time points, according to the 5$^{th}$ and 95$^{th}$ percentile over all frequencies for oscillatory event coincidence effects (correct > error, cluster permutation corrected p<0.05).

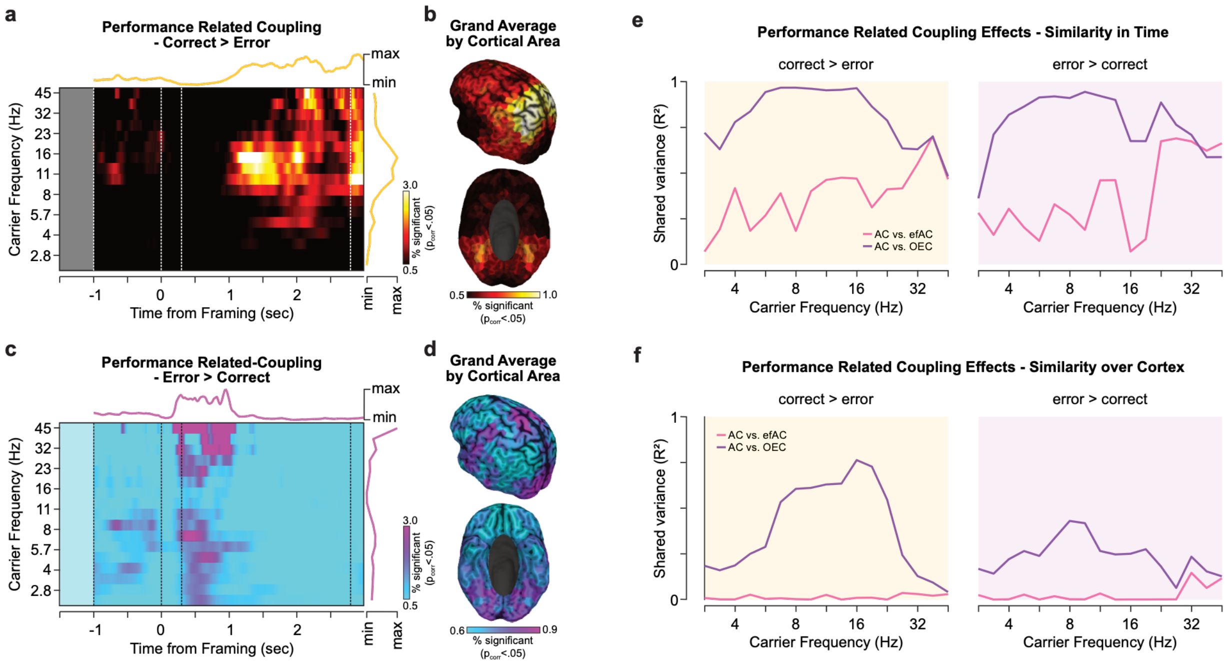


**Supplementary Figure S20 | Temporal, spectral and spatial distribution of performance related amplitude coupling.** The Figure extends the findings reported in Fig. 3 to full amplitude coupling (including high-amplitude events). **a** The time and frequency resolved distribution of performance related (correct > error, cluster permutation corrected $p<.025$) amplitude coupling connections. Colored lines on top and to the right denote the average over frequencies and time, respectively. **b** The average over the full time and frequency resolved distribution of behaviorally modulated amplitude coupling within each cortical source. **c-d** The same as **a-b** for the difference error > correct (uncorrected $p<.025$). **e-f** Pearson correlation of the behaviorally modulated (correct vs. error) coupling effects of amplitude coupling (AC) with event free amplitude coupling (pink line, efAC) and oscillatory event coincidence (purple, OEC) comparing the **e** temporal (see panel **a,c** and Fig. 3a,c) and **f** spatial (Supplementary Fig. S18 and S19) distribution of effects. The values depict the determination coefficient ($R^2$) within each frequency.

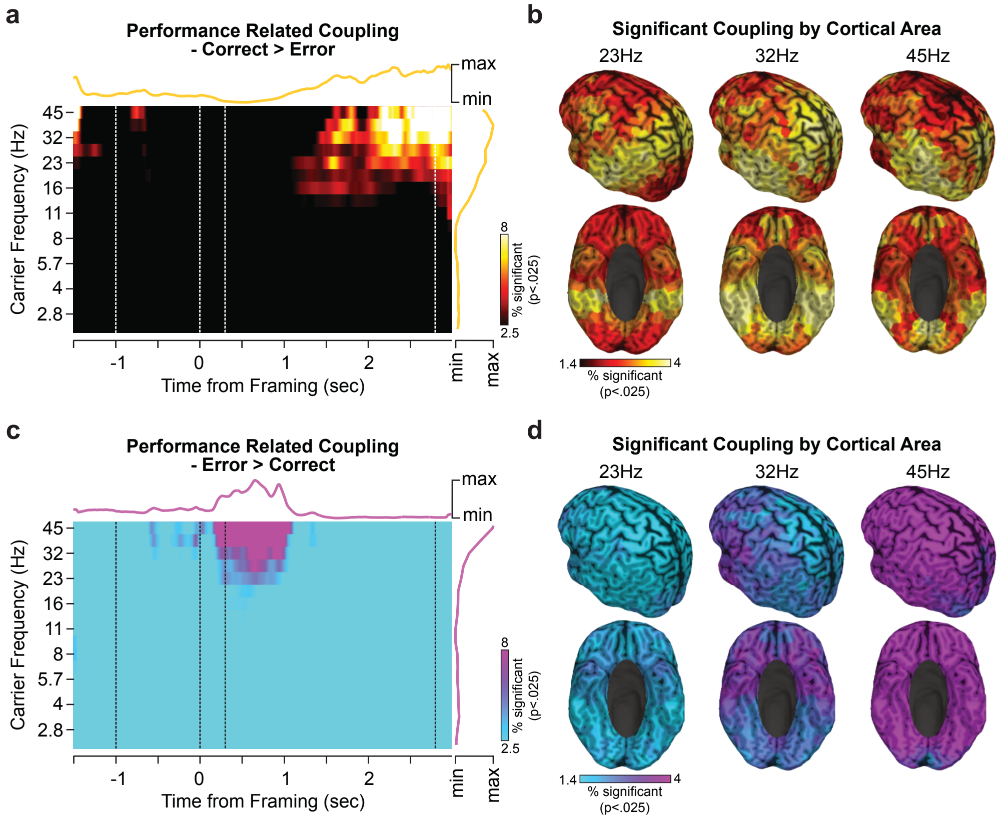


**Supplementary Figure S21 | Temporal, spectral and spatial distribution of performance related transient aperiodic amplitude coupling.** The Figure extends the findings reported in Fig. 3 to aperiodic event coincidences (AEC). **a** The time and frequency resolved distribution of performance (correct > error, one-sided t-test(19), uncorrected p<.025) aperiodic coupling connections. Colored lines on top and to the right denote the average over frequencies and time, respectively. **b** The time averaged frequency resolved distribution of behaviorally relevant aperiodic event coincidences within each cortical source for high-beta to gamma frequencies (23–45 Hz). **c-d** The same as **a-b** for the difference error > correct (one-sided t-test(19), uncorrected p<.025).

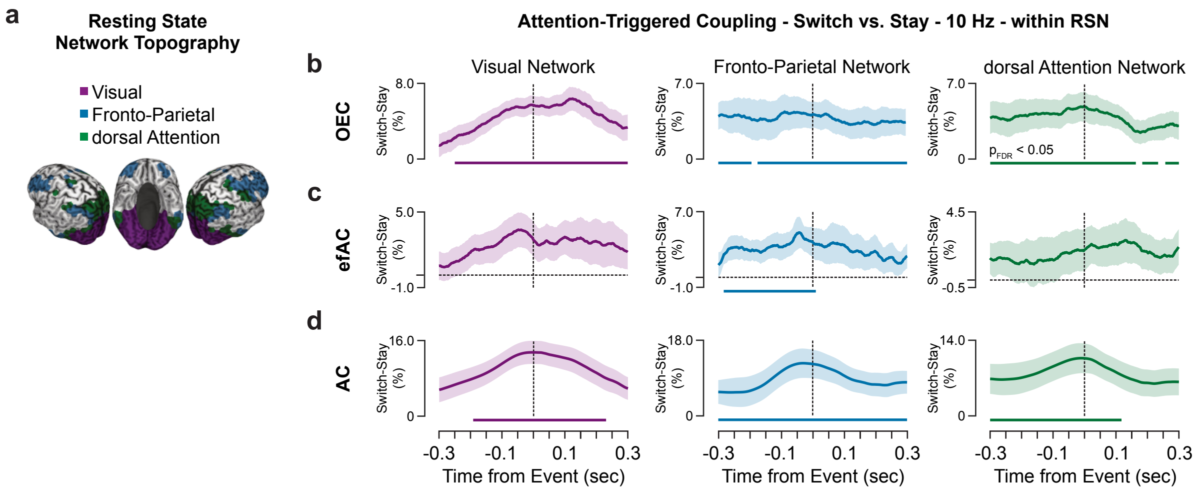


**Supplementary Figure S22 | Attention-triggered modulations in coupling within canonical resting-state networks. a** The cortical topography of sources associated with the visual (purple), fronto-parietal (blue) and dorsal attention (green) resting state network (RSN)[27]. **b-d** Difference in attention-triggered coupling (switch-stay, percent change) at 10 Hz averaged over connections within the visual, fronto-parietal and dorsal attention canonical resting state networks for **b** oscillatory event coincidences (OEC), **c** event free amplitude coupling (efAC) and **d** amplitude coupling (AC), color-coded as depicted in **a**. Solid lines and shaded areas denote the mean and standard error over participants (n = 20). Thick lines on the bottom display significant differences (paired t-test for stay vs. switch, df = 19, $p_{FDR} < 0.05$, FDR-corrected over time).

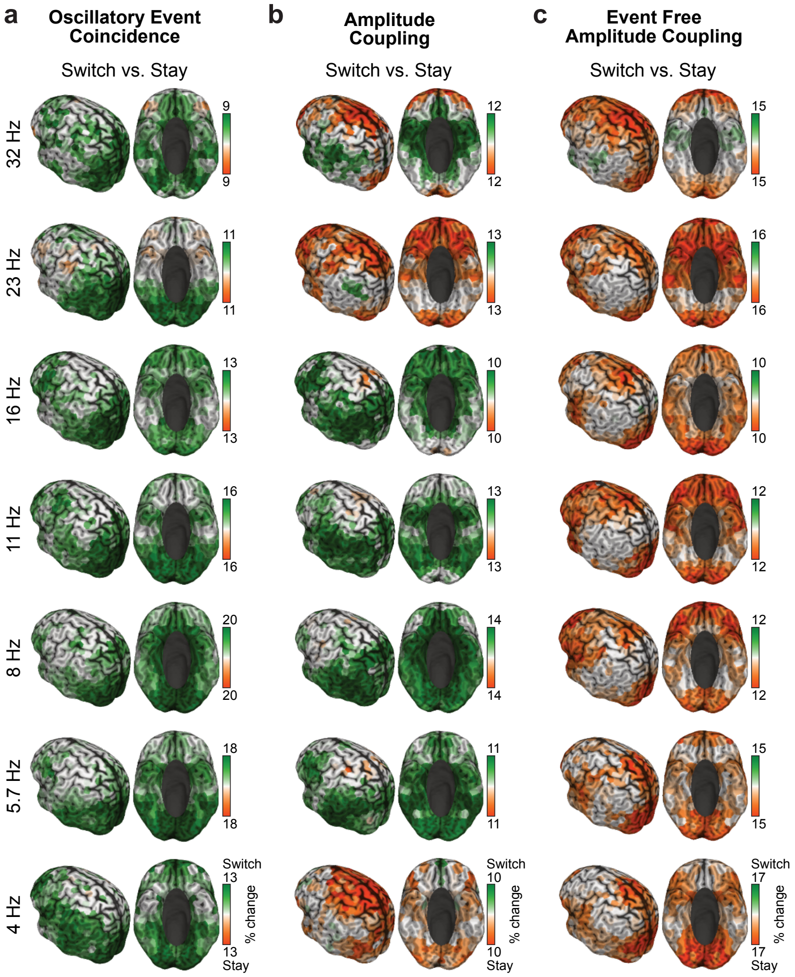


**Supplementary Figure S23 | Cortical distribution of attention modulated functional coupling.** Depicted are the percent change (modulation index) of attention-triggered (-0.05 to 0.05 s from switch or stay) coupling between saccade types, (switch-stay)/(switch+stay), for oscillatory event coincidence (left), amplitude coupling (middle) and event free amplitude coupling (right). The differences were averaged over all connections per source within a carrier frequency (4–32 Hz, bottom to top). The color scale was set to the 20$^{th}$ and 95$^{th}$ percentile of the absolute value of the difference within each panel.

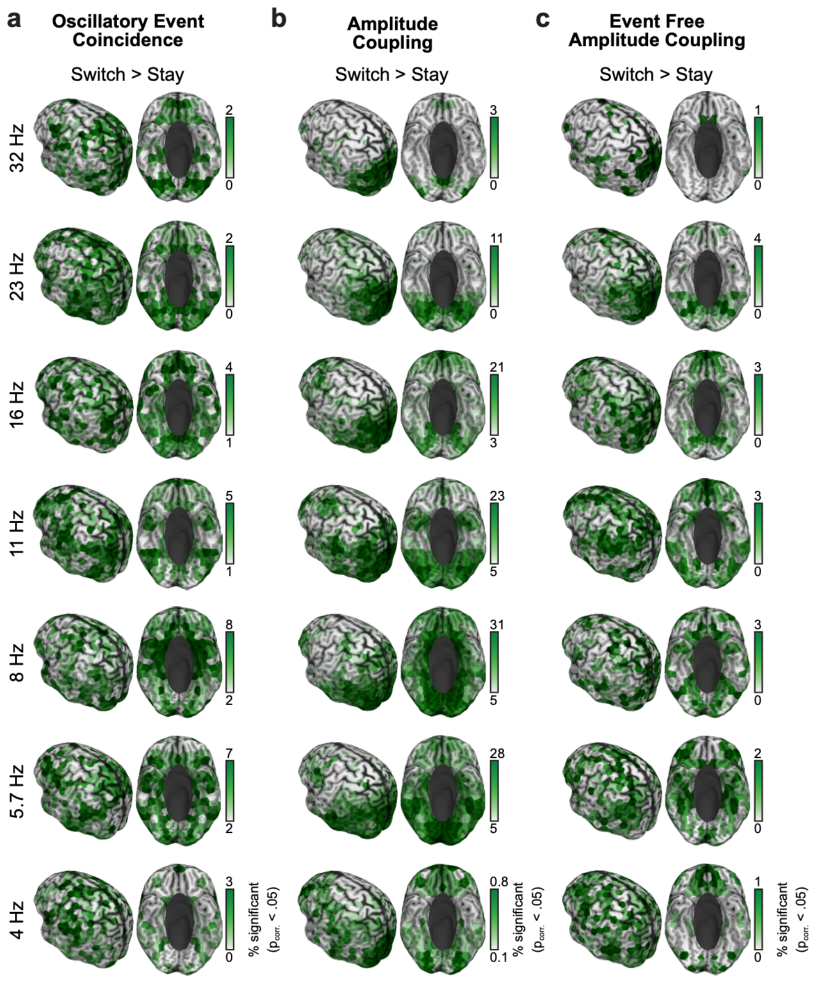


**Supplementary Figure S24 | Cortical distribution of functional coupling modulated during attention reallocation.** Depicted are the number of attention-triggered (-0.05 to 0.05 s from event) significantly modulated connections per source for oscillatory event coincidence (left), amplitude coupling (middle) and event free amplitude coupling (right). The coupling here was tested for switch > stay (paired $t_{switch>stay}$, df = 19, cluster permutation corrected p<0.05). The color scale was set between the $5^{th}$ and the $95^{th}$ percentile within each panel.

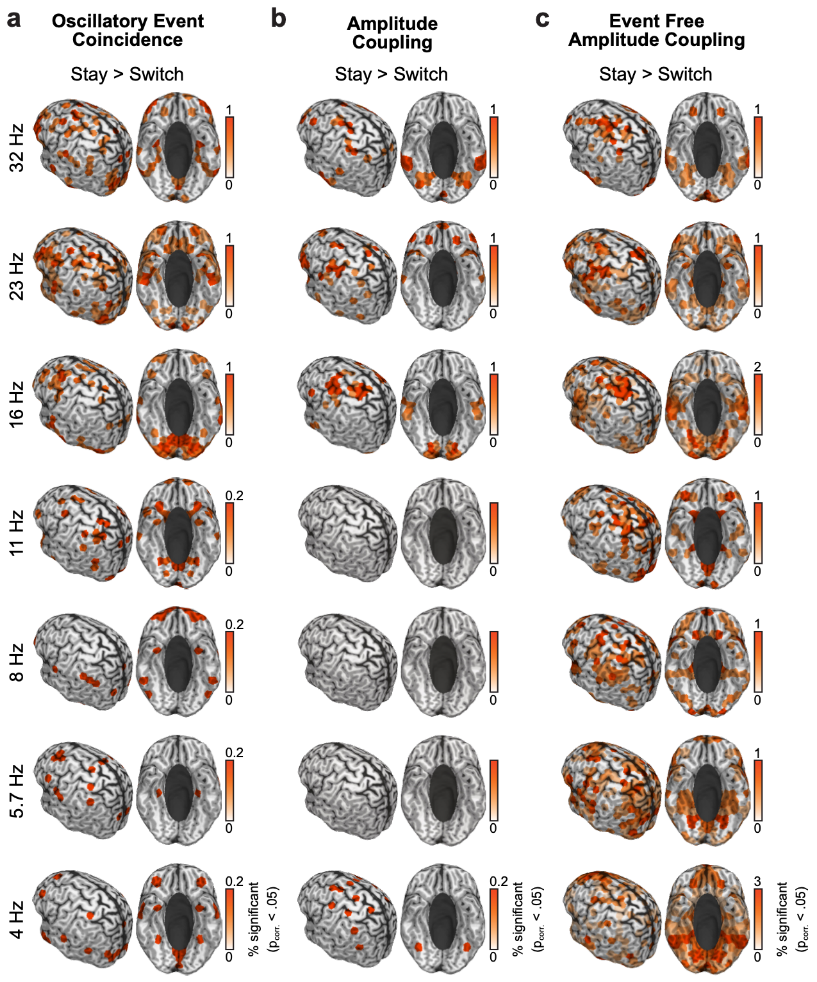


**Supplementary Figure S25 | Cortical distribution of functional coupling modulated during attention refocus.** Depicted are the number of attention-triggered (-0.05 to 0.05 s from event) significantly modulated connections per source for oscillatory event coincidence (left), amplitude coupling (middle) and event free amplitude coupling (right). The coupling here was tested for switch < stay (paired $t_{switch>stay}$, df = 19, cluster permutation corrected p<0.05). The color scale was set between the 5$^{th}$ and the 95$^{th}$ percentile within each panel.

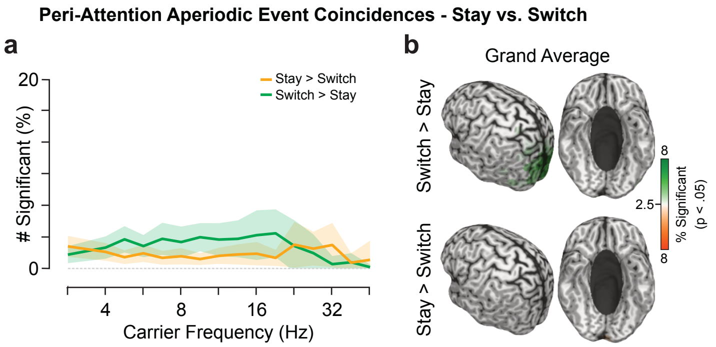


**Supplementary Figure S26 | Attention-modulated aperiodic event coincidences.** The Figure extends the findings reported in Fig. 4 to coincident high-amplitude aperiodic events (AEC). **a** Number of coupling connections modulated by attention (paired two-sided t-test(19), uncorrected p<.05) at the time of attention reallocation (switch) and refocusing (stay; averaged between -0.05 to 0.05 s) spectrally resolved between 2.8–45 Hz. Green and orange lines denote the mean proportion of connections per source increased during switches and stays, respectively. Shaded areas denote the standard deviation over cortical sources. **b** Grand average of the proportion of attention modulated coupling connections from each cortical source averaged over all frequencies. Color coded as above.

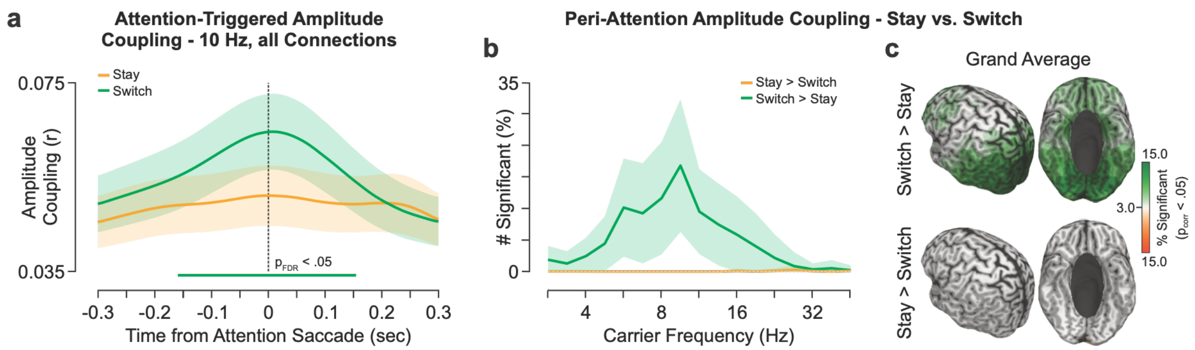


**Supplementary Figure S27 | Attention-modulated amplitude coupling.** The Figure extends the findings reported in Fig. 4 to full amplitude coupling (including events). **a** Attention-triggered amplitude coupling at 10 Hz separately for attention reallocation (switch, green) and refocus (stay, orange) averaged over all connections. Colored lines and shaded areas denote the mean and standard deviation over participants (n = 20). Thick lines denote the time of significant difference between switch and stay (two-sided, paired t-test, FDR-corrected over time, $p_{FDR}<.05$). **b** Number of amplitude coupling connections modulated by attention (paired two-sided t-test(19), cluster permutation corrected p<.05) at the time of attention reallocation (switch, green) and refocus (stay, orange; averaged between -0.05 to 0.05 s) spectrally resolved between 2.8–45 Hz. Green and orange lines denote the mean proportion of connections per source increased during switch and stay, respectively. Shaded areas denote the standard deviation over cortical sources. **c** Grand average of the proportion of attention modulated amplitude coupling connections from each cortical source averaged over all frequencies. The color scale is set between the $25^{th}$ and $95^{th}$ percentile separately based on the switch > stay (green) comparison.
